# An anti-amyloidogenic treatment to specifically block the consolidation of traumatic events in mouse

**DOI:** 10.1101/2020.01.21.913053

**Authors:** Paula López-García, Daniel Ramírez de Mingo, Kerry R. McGreevy, Anna Pallé, Helena Akiko Popiel, Andrea Santi, Yoshitaka Nagai, José Luís Trejo, Mariano Carrión-Vázquez

## Abstract

Post-traumatic stress disorder (PTSD) is a mental health disorder triggered by the exposure to a traumatic event that manifests with anguish, intrusive memories and negative mood changes. So far, there is no efficient treatment for PTSD other than symptomatic palliative care. Based on the implication of the functional amyloid cytoplasmic polyadenylation element binding protein-3 (CPEB3) in the consolidation of memory, we propose its active amyloid state as a possible therapeutic target by blocking the consolidation of traumatic memories through polyglutamine binding peptide 1 (QBP1), an inhibitor of the amyloid oligomerization previously investigated in *Drosophila*.

To test this idea in mammals, here we have developed a transgenic mouse that constitutively expresses QBP1 peptide. We first assessed the innocuousness of this peptide for the normal development of the animal, which also showed normal locomotor activity and anxiety. By performing a battery of standard memory paradigms, we then showed that hippocampal-dependent and aversive memories were impaired in the QBP1 mice. Furthermore, protein expression in the hippocampi of experienced mice showed that QBP1 mice do not increase their levels of amyloid oligomerization, evincing the blockade of the CPEB3 protein in its inactive state. The ability of QBP1 to block aversive memories in mice represents the proof of concept of a novel pharmacological approach for prophylaxis and therapy of acute stress and post-traumatic stress disorders.

## Introduction

<u>P</u>ost<u>t</u>raumatic <u>s</u>tress <u>d</u>isorder (PTSD) is a mental disorder classified within the group of disorders related to the exposure to traumas and stress factors in the Diagnostic and Statistical Manual of Mental Disorders 5 Approximately 7-8% of the population will experience PTSD at least once in their lives. Nowadays, <u>c</u>ognitive <u>b</u>ehavioral <u>t</u>herapy (CBT) is the main treatment for it while pharmacology is only used to treat the symptomatology^2–5^. One of the approaches to reduce the prevalence and severity of PTSD is to intervene in the psychological condition that occurs in response to exposure to trauma within the first month, the so-called <u>a</u>cute <u>s</u>tress <u>d</u>isorder (ASD)^6–8^. Currently, the specific pharmacology for the disease and its prophylaxis is based on promising evidence with hydrocortisone and propranolol^9,10^ as well as animal models based on stress responses and depression symptoms^11^. However, actually the limitation of the consolidation of traumatic memories has been the main objective of study in the design of a specific therapy^12–15^.

In this line, the discovery of prion-like domains in RNA regulatory proteins that are involved long-term memory^16–20^, unveiled an important mechanism for the stabilization of synaptic growth related to learning^21–25^. Thus, memory storage and underlying synaptic plasticity are mediated by the increase in the concentration of the <u>c</u>ytoplasmic <u>p</u>olyadenylation <u>e</u>lement <u>b</u>inding protein-3 (CPEB3) and its amyloid conversion^18,20,26^. In vertebrates, CPEB is usually coded by four genes with the first member to be identified being CPEB1, whereas CPEBs 2-4 were described later in mice^27–30^. Among these four vertebrate CPEB family proteins, there is a high similarity in its C-terminal domain (RNA-binding motif) while they vary widely in the N-terminal region (prion-like domain) with only CPEB2 and CPEB3 having regions rich in glutamine/asparagine residues (Q/N region), which resemble their orthologues in *Aplysia* and *Drosophila* (*Ap*CPEB and Orb2, respectively)^27^. On the other hand, it has been described the assembly of CPEB4 into liquid-like droplets and their implication on cell cycle regulation^31^, although its function seems to be dispensable for memory consolidation^32^.

Experiments in yeast and in culture of sensory neurons from *Aplysia* have revealed that *Ap*CPEB is a functional prionoid whose mechanism of conversion to the transmissible state is involved in the long-term potentiation of *Aplysia’s* motor and sensory synapses^17,22,33,34^. It has also been shown that Orb2 is necessary for the maintenance of long-term memory through a mechanism that requires its N-terminal prion-like domain^24,35–37^, and only those neurons in which Orb2 oligomers are deposited are able to storage long-term memory in *Drosophila*^38,39^. Despite the lowering in the number of glutamine residues in the polyQ tracts observed in the prion-like domain of the CPEB family throughout evolution, a similar functionality of the amyloid oligomers is assumed in the mammalian CPEB^30,40^.

In this sense, the mammalian CPEB3 is able to form self-propagating and heritable aggregates in yeast and its aggregation in the mouse brain occurs only under physiological conditions after neuronal stimulation and is associated with LTM^41,42^. Furthermore, it is known that CPEB3 <u>k</u>nock <u>o</u>ut mice (KO) shows defects in memory consolidation associated with spatial learning and that the interruption in CPEB3 function after consolidation leads to an impairment in long-term memory^40,43^. Moreover, it also has been described that, while CPEB4 seems to be dispensable for hippocampus-dependent learning as mentioned, CPEB2 fulfills a similar function as CPEB3 and it is also required for long term memory consolidation^32,44^.

Glutamine-rich regions (polyQ) have been widely related to the formation of aberrant amyloids in pathologies such as Huntington’s disease and various types of spinocerebellar ataxia^19,45–47^. Nagai *et al.* (2000) developed the QBP1 peptide (sequence: SNWKWWPGIFD) that selectively binds to long *(i.e.,* pathological) polyQ stretches. Its therapeutic importance lies in its specific binding to the monomer of the protein and its ability to block the transition to the toxic conformation and thus preventing the amyloidogenic cascade at its start^48,49^.

Later, it was shown that QBP1 has some polyvalence as it is able to block the amyloidogenesis of other amyloids^50^. Recently, Hervás *et al.* (2016) have demonstrated that QBP1 inhibits *in vitro* the aggregation and amyloidogenesis of the Orb2 prion-like domain and blocks the consolidation of memory and learning in *Drosophila*^39,50^. Furthermore, it has also been shown that QBP1 blocks the amyloidogenesis of *Ap*CPEB *in vitro* (Hervás, R., Fernández-Ramírez, M., Galera-Prat, A., Suzuki, M., Y.N., Bruix, M., Menéndez, M., Laurents, DV. & M.C.V. Memory consolidation systems based on prion-like CPEB proteins show different amyloidogenic structural properties. *Submitted).* Thus, QBP1 works in these two lower organisms and the experiments in *Drosophila* indicated that chronic QBP1 expression impairs the formation of <u>l</u>ong-<u>t</u>erm <u>m</u>emories (LTM) while does not interfere in the formation of <u>s</u>hort-<u>t</u>erm <u>m</u>emory (STM).

In the present study, we have produced and characterized a constitutive QBP1 transgenic mouse with the purpose of implementing an anti-amyloidogenic approach for the blockade of the oligomerization of CPEB3, so that we could effectively block the long-term consolidation of new events, including aversive ones. Our results represent a proof of concept of the validity of a specific pharmacological approach for preventing the development of PTSD and ASD based on the blockade of memory consolidation of traumatic events.

## Results

### QBP1 transgenic mice have normal neurological, locomotor activity and anxiety

Previous results have shown that QBP1 blocks *in vitro* human CPEB3^51^, which sequence identity with the mouse orthologue is 97% in the N-terminal region^52^. To investigate the effect in memory consolidation of QBP1 in mouse CPEB3 (mCPEB3) we followed a similar approach to the one previously used in *Drosophila*^39^, *i.e.* we generated a transgenic mouse expressing two copies of QBP1 in tandem fused to the <u>g</u>reen <u>f</u>luorescence <u>p</u>rotein (eGFP) with its transcription regulated by the CAG mammalian promoter for its constitutive expression **[Figure 1A]**. To test the correct presence of the peptide into the three QBP1 constitutive transgenic lines generated by Japan SLC, brain lysates from one mouse of each line were tested by western blot analysis. We detected QBP1 only in one of these lines **[Figure 1B]**, which was used to produce all the QBP1 transgenic mice used in these experiments. Moreover, the immunohistochemical analysis confirmed the presence of QBP1 in the hippocampus **[Figure 1C]**.

**Figure 1:**
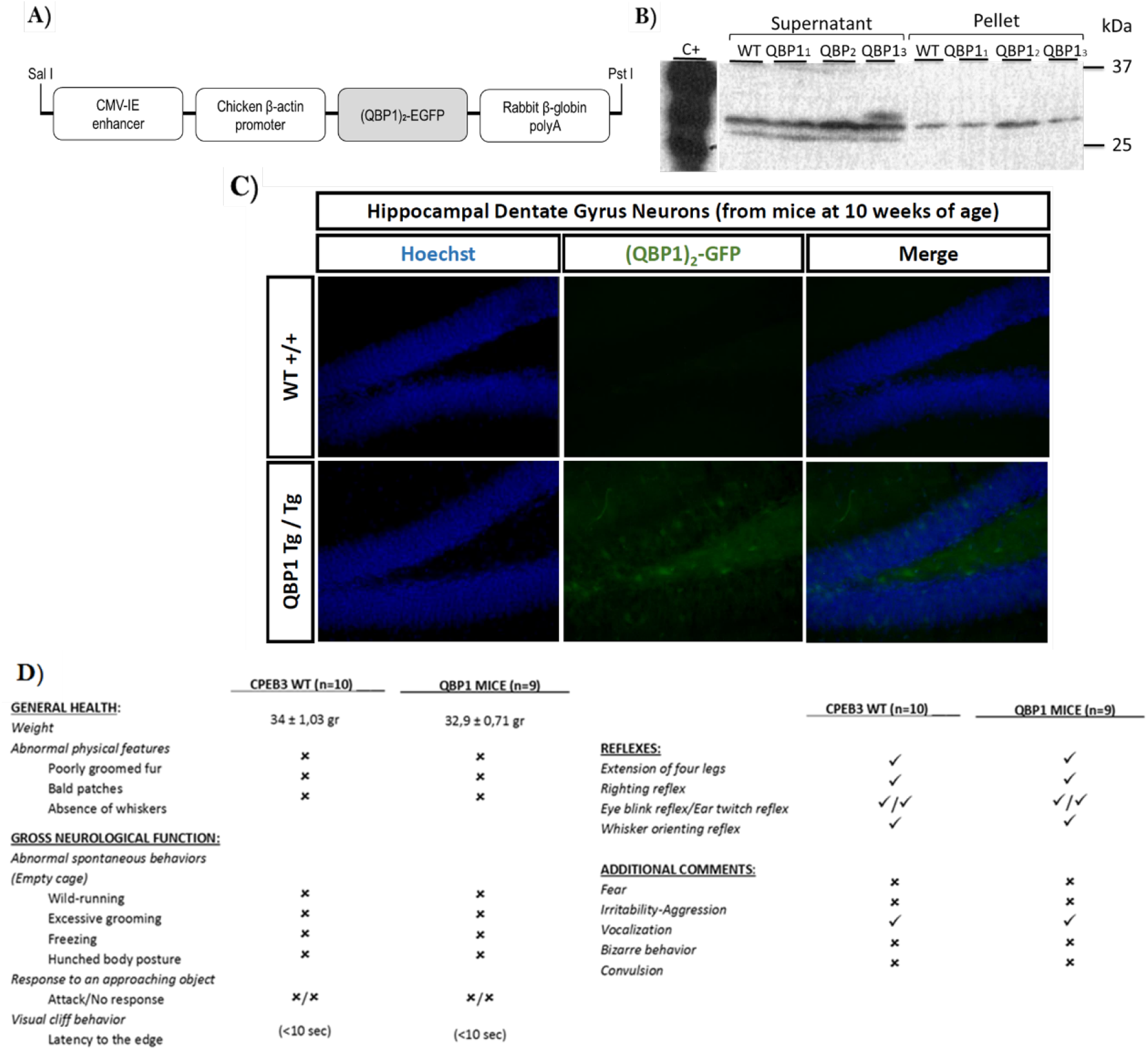
QBP1 transgenic mouse: generation, characterization and QBP1 expression. **A) Scheme of the construct used to generate the QBP1 transgenic mouse.** This construct was used to microinject fertile eggs into pseudopregnant females, using the CAG promoter for its constitutive expression in mammals and long maintenance of protein expression. **B) Western blot of the generated transgenic lines**. Of the three lines produced, only in one of them (QBP1-3) it was confirmed the expression of the peptide in the supernatant and was the one used to generate the breeding mouse. Q19-YFP expressing COS-7 lysate was used as a positive control, revealed with an anti-GFP (1:500) and a secondary mouse IgG-HRP (1:2000). **C) Confocal images of hippocampal sections** obtained from WT and QBP1 transgenic mice, in which EGFP was observed because of its natural fluorescence and Hoechst was used as a cell viability control. DNA counterstain for nuclei. Scale bar: 50 μm. **D) General health characterization** of QBP1 transgenic mouse according to (Crawley, 1999)^61^.

We started the phenotype characterization of this new transgenic mouse with established and reproducible behavioral tasks of general health, in order to discard gross defects. We measured body weight throughout the course of the experiments, corroborating a normal increase with time both in wild wype (WT) littermates and QBP1 transgenic mice [F_WEIGHT(5,13)_=42.174 p<0.001], with no differences between them [F_INTERACTION(5,13)_=0.972 p=0.470]. Furthermore, the transgenic mouse showed normal gross neurological functions (wild-running, visual cliff, approaching object) as well as normal grooming and walking body posture **[Figure 1D]**.

The characterization of the transgenic mouse was carried out following the scheme shown in **Figure 2A**, which follows an order that minimizes the impact on the mouse behavior. In comparison with WT littermates, QBP1 mice showed normal locomotor activity in the **activity cage** (two-day protocol): in the horizontal activity **[Figure 2B-i]**, which measures the number of times that mouse interrupts the horizontal sensor, there were no significant differences between groups and which varies with time in day 1, thus establishing that the locomotor activity of QBP1 mice was normal [F_DAY_1(4,68)_=27.841 p<0.001 and F_INTERACTION(4,68)_=1.793 p=0.140; inter-subject effect F_DAY_1(1,17)_=1.124 p=0.304 and F_INTERACTION(1,17)_=620.453 p<0.001)]. In day 2, which analyzes the mouse activity in a known environment, QBP1 mice showed an abnormal and significant increase in their total horizontal activity (p=0.001). Furthermore, analyzing this activity *per* minute **[Figure 2B-ii],** WT mice maintained a linear activity that did not vary with time while the QBP1 mice showed an unusual activity for a known environment [F_DAY_2(4,60)_=4.665 p=0.002 and F_INTERACTION(4,60)_=2.268 p=0.072; inter-subject effect F_DAY_2(1,15)_=17.819 p=0.001 and F_INTERACTION(1,15)_=268.620 p<0.001], increasing their activity as if they are re-exploring the cage [F_QBP1(4,12)_=4.751; p=0.016]. Moreover, this was reflected in the differences between groups that exist in the last minutes of day 2 of the protocol: minute 3 (p<0.001), minute 4 (p=0.002) and minute 5 (p= 0.007).

**Figure 2:**
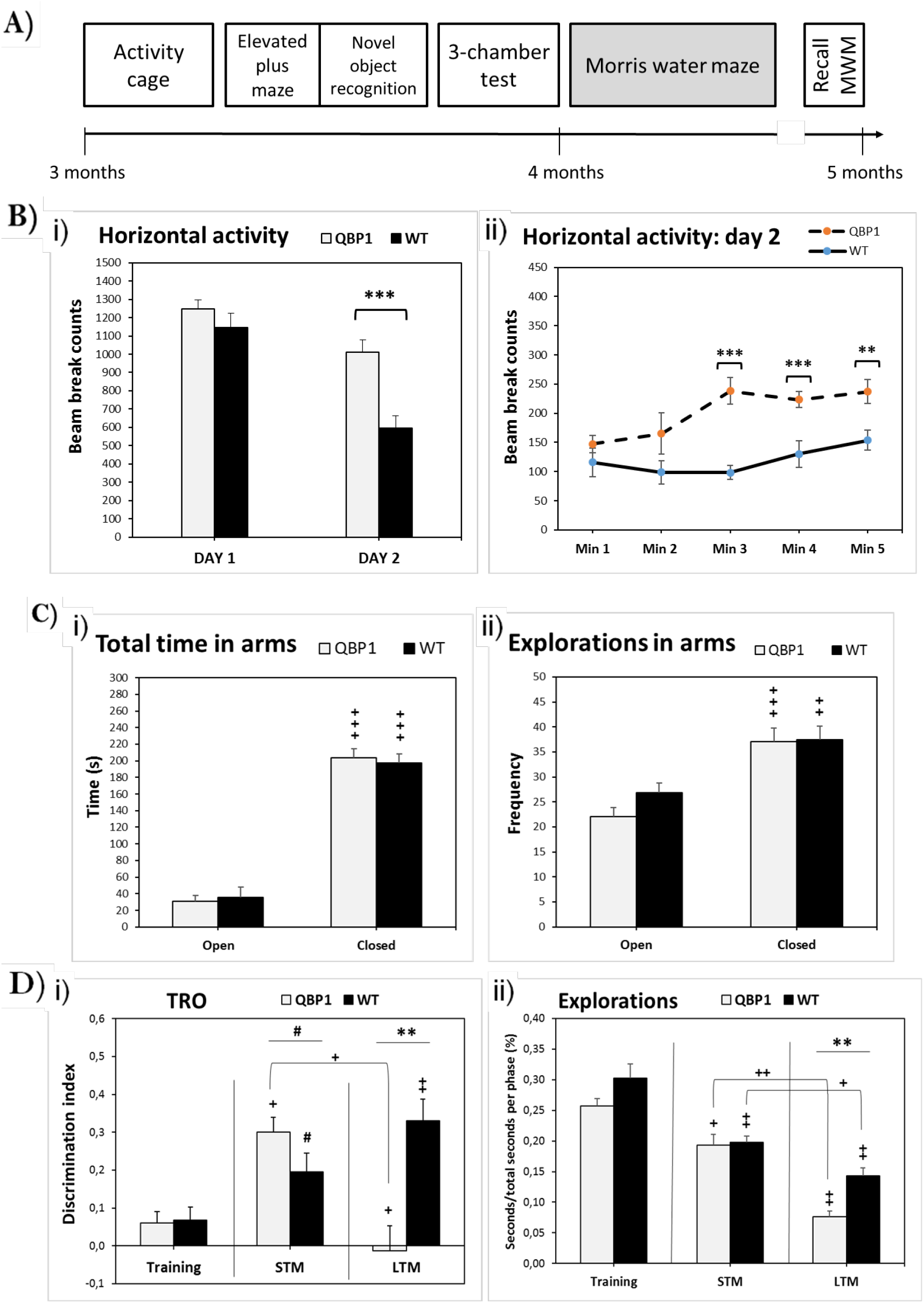
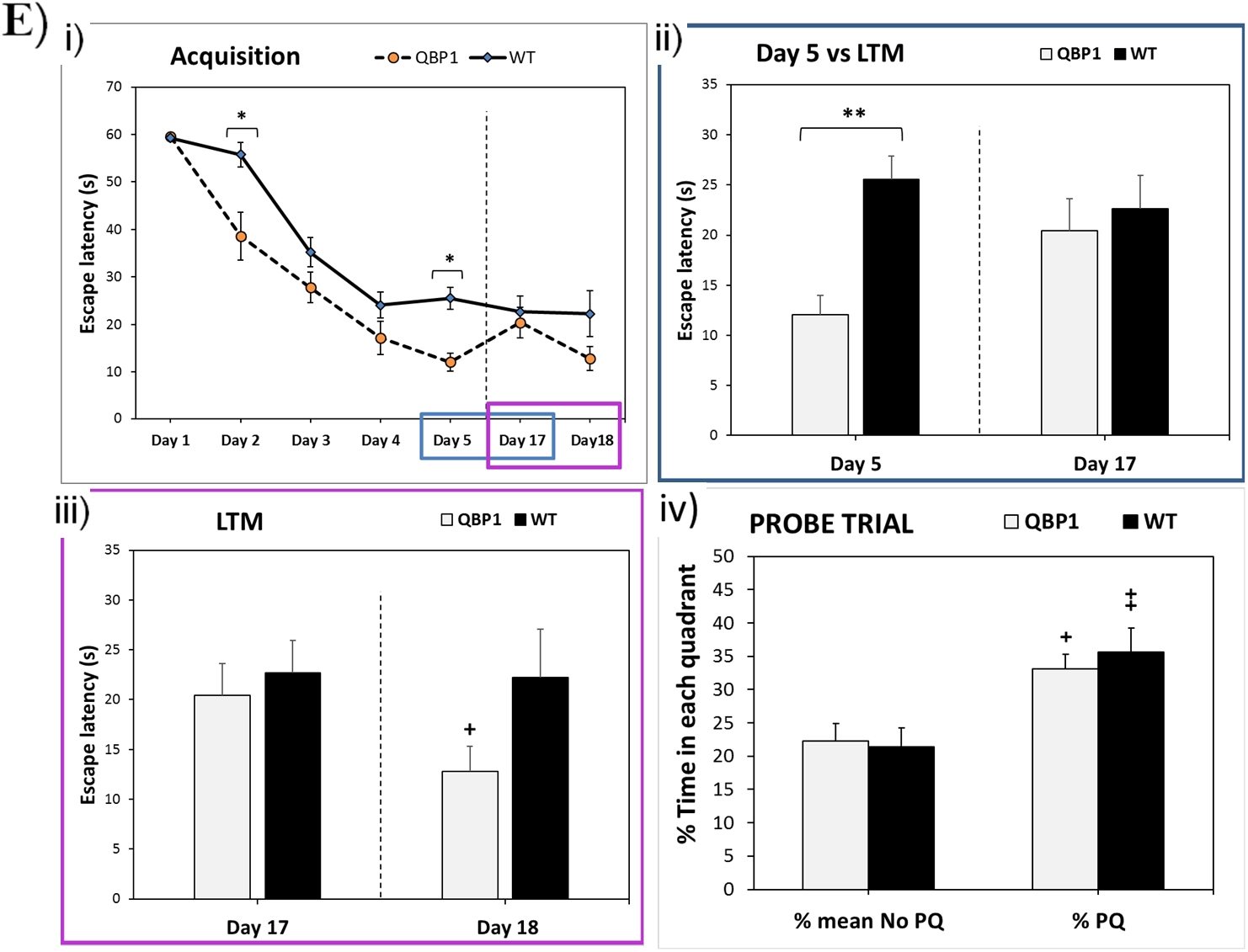
Analysis of hippocampal dependent memories in the QBP1 transgenic mouse. **A) Scheme of the test temporal sequence implemented in the QBP1 transgenic mouse**. **B) Results of locomotor activity of the QBP1 mice in a two-day protocol**: the first day is meant to assess locomotor activity in a novel environment, while the second day is meant to analyze the locomotor activity in a known environment. **i)** Total horizontal activity of both days of the protocol, in which at day 1 there are no differences and at day 2 QBP1 significantly increases its activity in comparison with control mice (p=0.001). **ii)** Horizontal activity per minute of day 2, where the activity of QBP1 mice significantly increases in the last minutes of the test (p<0.001; p=0.002 and p=0.007 respectively). **C) QBP1 mice exhibit a normal anxiety behavior in the Elevated Plus Maze test**. There are no significant differences among groups in any of the variables analyzed (explorations and total time). **i)** In the total time, QBP1 mouse spends very significantly longer time in the closed arms than in the open arms (p<0.001). **ii)** At the number of explorations, WT and QBP1 mice had a higher significance at explorations in closed arms in comparison with open arms (p<0.001for QBP1 and p=0.003 for WT). **D) Long-term memory is impaired in Novel Object Recognition test**. **i)** The DI values in each phase test are showed. WT mice tended to show higher DI scores in the STM phase (p=0.059) and significantly higher DI scores in the LTM phase (p=0.007). QBP1 mice showed significantly higher DI scores at STM phase compared to the training phase (p=0.01), but they showed significantly lower DI scores in the LTM phase (p=0.025) and they significantly reduced their DI scores in the LTM phase compared to the STM phase (p=0.012). Finally, QBP1 mice tended to show higher DI scores than WT animals in the STM phase (p=0.057) and significantly lower DI scores in the LTM phase (p=0.002). **ii)** At explorations, the number are reduced as object’s exposure is increased (p=0.008 with STM and p=0.005 with LTM for WT; p=0,017 with LTM and p=0.008 for QBP1), and only there are differences between subjects in the LTM phase (p=0.001). **E) Long-term memory is altered in the Morris Water Maze test**. **i)** Acquisition of the platform location: escape latency of mice shows a better performance in the QBP1 mice and significance difference between groups in day 2 (p<0.001) and day 5 (p<0.001). **ii)** Comparison of the latency of escape between day 5 of acquisition and day 17 (LTM), which reflects a loss of the significance of the last day of acquisition with respect to the long-term test in the QBP1 animals. **iii)** Comparison of the latency of escape between day 17 and day 18 (LTM), which reflects an improvement of the escape latency in the QBP1 mice (p=0.04) 24h after the LTM test. **iv)** Time percentage in each quadrant in probe trial (probe Q), in which both groups spend significantly more time in PQ than in the other quadrants mean (p=0.004 in WT and p=0.014 in QBP1). A proportions binomial distribution was used to analyze this comparison. Data were analyzed by a mixed model ANOVA, Friedman test, or repeated measures ANOVA (for MWM), using T test, Mann Whitney *U*, and Wilcoxon signed-rank test to assess intra- and intersubject differences (showing mean ± SEM). For comparisons between independent groups: *p<0.05, **p<0.01, ***p<0.001, trends 0.05 ≥ #p < 0.09; for comparisons between dependent groups: +p<0.05, ++p<0.01, +++p<0.001, trends 0.05 ≥ #’p < 0.09.

The characterization of the QBP1 mouse also included an **<u>E</u>levated <u>P</u>lus <u>M</u>aze test (EPM)** to evaluate anxiety. We did not find significant differences in any of the variables analyzed (number of entries, explorations and total time spent in the open and closed arms), which suggests normal levels of anxiety in QBP1 mice **[Figure 2C-i, *entries data not shown*]**. Both groups of animals spent significantly more time in the closed arms of the maze compared to the time spent in the open arms (p<0.001). Regarding the number of explorations **[Figure 2C-ii]**, both groups explored the closed arms of the maze significantly more compared to the open arms of the maze (p=0.003, WT and p<0.001, QBP1).

To assess the normal sociability of the transgenic mouse, a **3-chamber test** was used to evaluate social preference and social novelty preference ***[supplementary material, Figure S1*]**. <u>On the first day</u> a sociability study is carried out in which the ethological measure of the time they spend in a social compartment (confined with a stranger male mouse) *versus* an empty one is analyzed. We found no differences in the time they spent exploring the social compartment, as both displayed a preference for the social one. Thus, both groups displayed normal sociability-like behavior. This was also confirmed using a ratio for social preference [***Figure S1-D]*** in which no differences between groups were observed. However, we found a trend for a higher number of entries into the social compartment in the QBP1 group [***Figure S1-C*]**. On <u>the second day</u>, when social memory was tested, we found no differences between groups in the percentage spent in each chamber, as both groups spent the same time exploring a compartment with a stranger novel mouse compared to that with a known mouse [***Figure S1-E*]**. Furthermore, no differences were found in the ratio of preference [***Figure S1-G*]**, indicating that none of the groups displays a preference for a novel mouse. Additionally, on day 2 we found that overall WT mice performed more entries independently of the compartment [***Figure S1-F*]**.

### Long-term memory of the QBP1 mice is impaired

To assess our working hypothesis that the anti-amyloidogenic QBP1 peptide expressed in mice would interfere with the oligomerization of mCPEB3 and block memory consolidation in turn, firstly, a **<u>N</u>ovel <u>O</u>bject <u>R</u>ecognition test (NOR)** was performed. It evaluates rodents’ ability to recognize a novel object in an environment in the absence of reinforcements **[Figure 2D]**. When rodents are exposed to a familiar and novel object, they spend more time exploring the novel object as they have a natural preference for novelty. According to the <u>d</u>iscrimination <u>i</u>ndex (DI),both groups significantly varied their performance throughout the test (p=0.003, WT and p=0.001, QBP1; Friedman Test) **[Figure 2D-i]**. *Post hoc* analysis indicated that WT animals tend to show higher DI scores in the STM phase (p=0.059), and significantly higher DI scores in the LTM phase (p=0.007), compared to the training phase. These results suggest that WT animals were able to discriminate the novel object, especially in the LTM phase.

On the other hand, QBP1 mice were able to discriminate the novel object in the STM compared to the training phase (p=0.01), but they were unable to discriminate the novel object in the LTM phase. QBP1 mice showed significantly lower DI scores in the LTM phase, compared to the training phase (p=0.025) and they significantly reduced their DI scores in the LTM phase compared to the STM phase (p=0.012). Furthermore, QBP1 mice tend to show higher DI scores than WT animals in the STM phase, while they showed significantly lower DI scores in the LTM phase (p=0.057 and p=0.002, respectively). Taking together, these results suggest that although QBP1 mice were able to discriminate the novel object 1 hour after training, they were unable to discriminate the change of object after 24 hours, indicating a LTM impairment.

Regarding exploration, both groups of mice significantly reduced their exploration time throughout the test (p<0.001; Friedman Test), which is normal because they have already been exposed before to the task and their discrimination becomes easier since they have learn. **[Figure 2D-ii]**. *Post hoc* analysis indicated that WT mice reduced their exploration time in the STM phase and the LTM phase, compared to the training phase (p=0.008 and p=0.005, respectively) and they significantly explored less in the LTM phase compared to the STM phase (p=0.016). QBP1 mice also significantly reduced their exploration time in the STM phase and the LTM phase, compared to the training phase (p=0.017 and p=0.008, respectively) and they significantly explored less in the LTM phase compared to the STM phase (p=0.008). QBP1 mice explored significantly less than WT animals in the LTM phase (p=0.001), which was the phase where the QBP1 animals were unable to discriminate the novel object.

Memory consolidation of QBP1 mice was also tested by the **<u>M</u>orris <u>W</u>ater <u>M</u>aze (MWM)** paradigm, a hippocampal-dependent task widely used to study spatial learning and memory in rodents. In the acquisition phase, QBP1 mice and the littermates significantly decreased their escape latency along days, which showed that they have learned the task [F_DAYS(4,68)_=83.342 p<0.001 and F_INTERACTION(4,68)_=3.050 p=0.023]. The inter-subject effect reflected that QBP1 mice seemed to performed significantly better than WT mice on each day [F_DAYS(1,17)_=15.854 p=0.001 and F_INTERACTION(1,17)_=1002.673 p<0.001)]. Furthermore, QBP1 mice showed lower escape latency than that of WT mice (p<0.001) on the last day of acquisition (Day 5), probably because WT mice memory reach a plateau on that final day **[Figure 2E-i]**.

In view of these differences in the acquisition days along trials, we decided to perform an <u>i</u>ntertrial <u>a</u>cquisition <u>r</u>ate (**IAR) index**, which revealed the differences in mouse performance in the first trial compared to the mean within a single day [***Figure S2-A*]**. During the first acquisition days, the QBP1 mouse had an increased index (~10), which means that after resting overnight their performance was worse in the first trial, but in the last trial of each day -*after four repeated exposures*-their performance was much better. Finally, QBP1 mice maintained an index of ~ 0 on day 4-5, which indicates that they have improved their performance in the first trial.

However, when mice are exposed to the task again 12 days later to explore their long-term memory (called Day 17), QBP1 mice increased the escape latency in comparison to Day 5 **[Figure 2E-ii]**. This increase represents a substantially worse performance of the transgenic mouse at longterm on day 17 in comparison to their performance on day 5, showing an intra-subject significance (p=0.047). The repetition of the test on day 18 resulted in a new decrease in the escape latency of the QBP1 mouse compared to the previous day, as its performance was even better than their littermates on that day. Although there were no significant differences between groups as in day 5, there was a new significant difference in QBP1 mice between day 17 and 18 (p=0.043), which suggests a substantial improvement explained by the re-exposition to the test the day before **[Figure 2E-iii]**.

We also performed a <u>probe trial</u> 24h after acquisition and we found no differences between both groups, as they both spent more time in the quadrant (probe Q) where the platform was **[Figure 2E-iv]**. Furthermore, we analyzed the <u>probe P</u> to estimate at the millimeter level the virtual location of mouse search in the platform [***Figure S2-B*].** Only WT mice crossed significantly more times the virtual platform location in the correct position instead of the same location in the others quadrants (p=0.017), suggesting that only WT mice retained the complete spatial knowledge of the task with time.

### QBP1 impairs long-term consolidation of aversive memories

To analyze the effect of the QBP1 on aversive memories, we performed the implantation of a traumatic memory (inescapable electric shock) associated to a context (conditioned stimulus)^53^. The **<u>c</u>ontextual <u>f</u>ear <u>c</u>onditioning (CFC)** was tested following the scheme showed in **Figure 3A**. An **Open field test** was performed to ensure there were no differences between groups in mouse activity and anxiety before the fear implantation. The results obtained showed that both groups have a similar activity pattern as shown by the total distance traveled over test duration **[Figure 3B-i]**, which confirmed that the transgenic mouse has normal locomotor activity. The analysis of the anxiety behavior provided by this test showed that there were no significant differences between groups (p=0.106), both spending more time at the margin compared to the center (p<0.001), replicating the results observed in the Elevated Plus Maze **[Figure 3B-ii]**.

**Figure 3:**
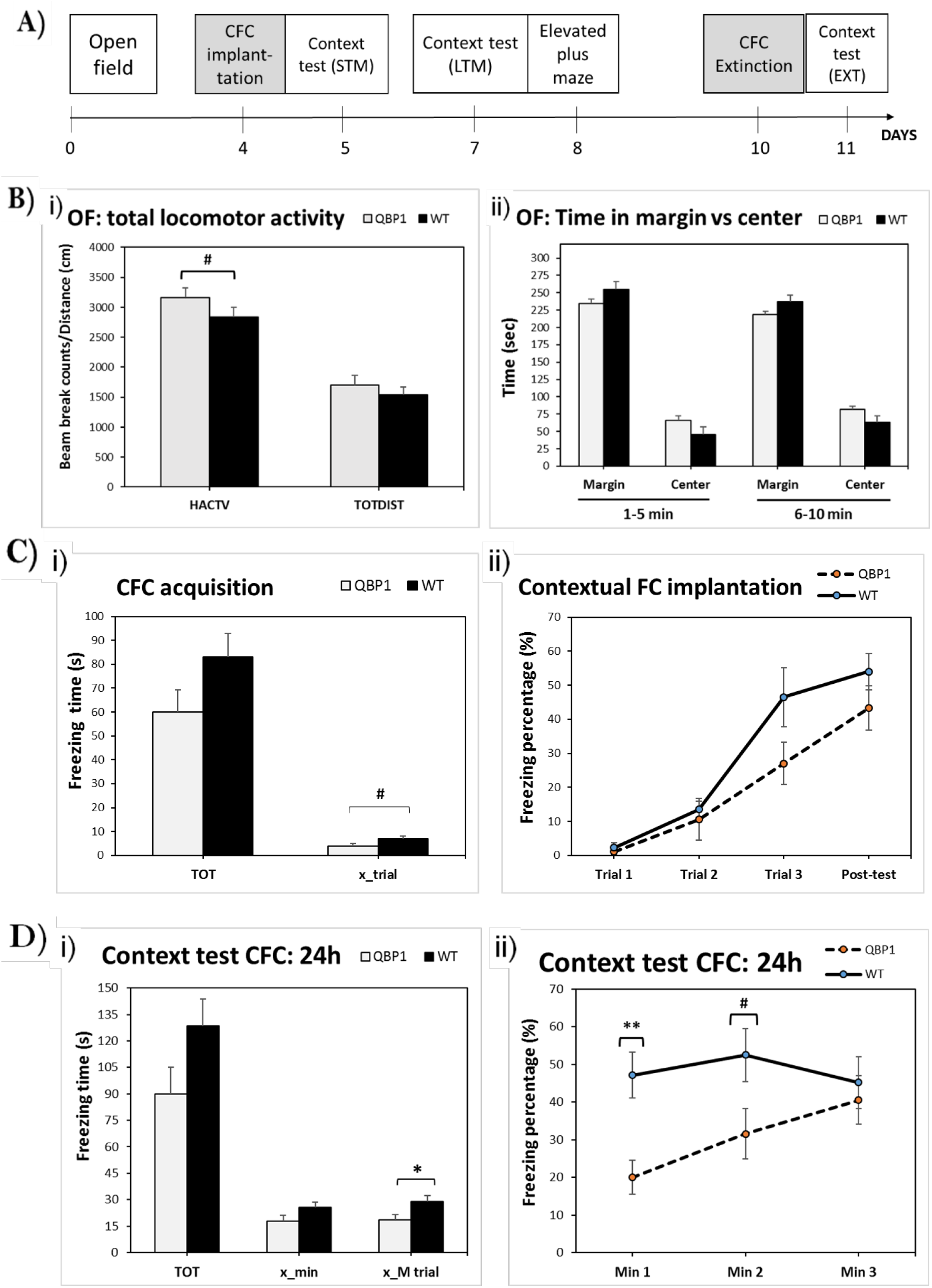
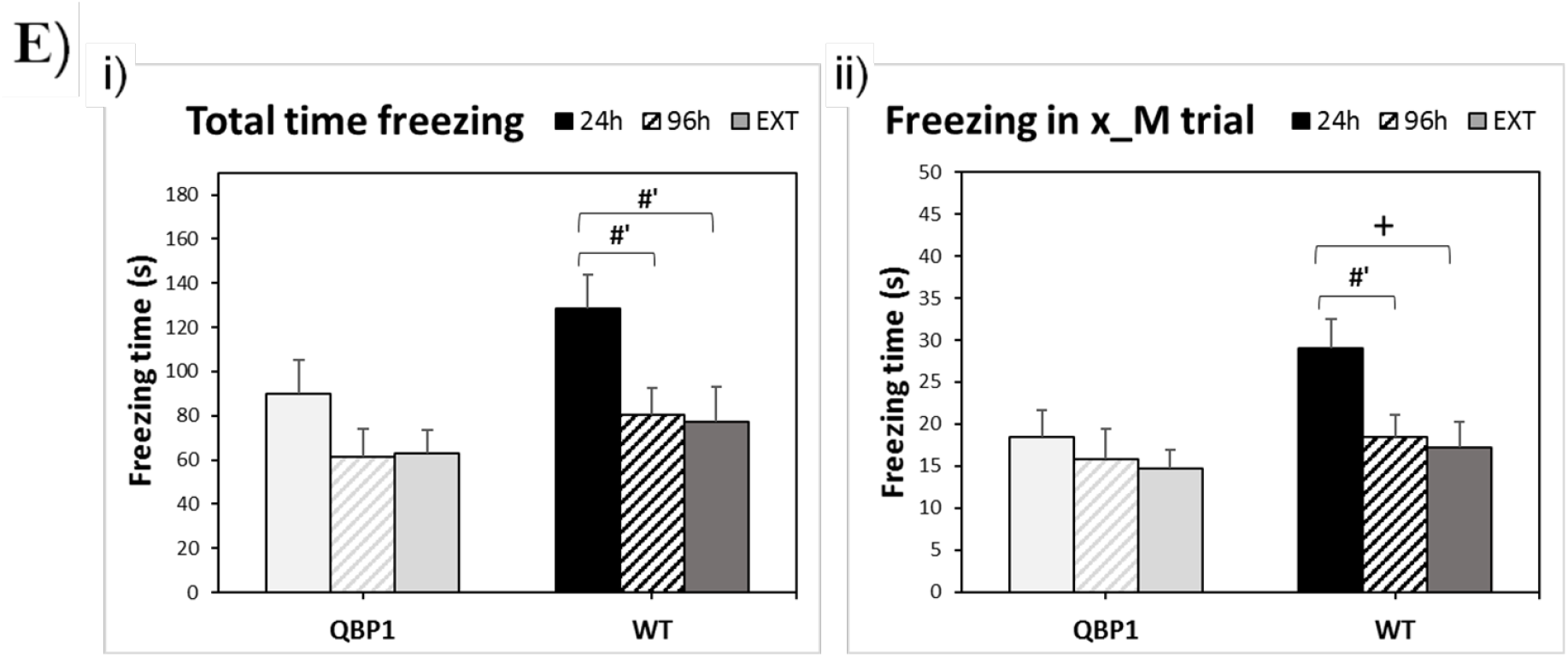
Analysis of aversive memories in the QBP1 transgenic mouse. **A) Diagram of the test temporal sequence implemented in the QBP1 transgenic mouse for the fear conditioning implantation**. **B) Open field showed normal anxiety level in the QBP1 transgenic mouse**. **i)** Total horizontal activity and total distance traveled (total time) showed no differences between groups, reflecting just a trend in total horizontal activity (p=0.057). **ii)** Time spent in the margin *versus* the center showed that QBP1 and WT mice spent more time in the margin than in the center (p<0.001), without any differences between groups (p=0.106). **C) Contextual fear conditioning was implanted in the QBP1 mice showing that aversive memories are impaired**. **i)** Freezing total time (TOT) and the average freezing time in trials segments *(x_trial)* at fear acquisition is showed. WT mice showed a trend to increase their *x_trial* (p=0.076) in comparison to QBP1 mice. **ii)** The acquisition phase analyzed by the freezing percentage on each trial segment showed that both groups are able to learn the fear conditioning time as they increase its freezing time among trials, without significant differences between them. **D) Fear learning is reflected in a context test 24 after the acquisition**. **i)** Freezing total time (TOT) and average freezing time per minute at context test *(x_min)* reflected that there were any differences between groups. The average freezing time in Min 1-2-3 *(x_M trial)* is also represented, in which WT mice significantly increased their freezing time (p=0.048). **ii)** Freezing percentage on the representative test minutes is shown, reflecting a significant difference at the freezing percentage in Min1 between WT mice and QBP1 mice (p=0.005), which correspond to the first trial of the acquisition phase of fear conditioning, and a trend in Min2 (p=0.055). **E) Intra-subject comparison of freezing time on each context test. i)** Total freezing time showed no differences along time and test in the QBP1 mice. Unlike, WT mice reflected a trend to decrease their freezing at 96h (p=0.081) and at extinction test (EXT, p=0.056) in comparison to their values at 24h. **ii)** The average freezing time in Min 1-2-3 *(x_M trial)* is represented for QBP1 mice and control littermates. Only WT mice showed differences along tests: there was a trend to decrease their freezing at 96h (p=0.071) and a significant decrease at extinction test (EXT, p=0.039) in comparison to 24h. Data were analyzed by a mixed model ANOVA, Friedman test or repeated measures ANOVA; using T test, Mann Whitney *U*, and Wilcoxon signed-rank test to assess intra- and intersubject differences (showing mean ± SEM). For comparisons between independent groups: *p<0.05, **p<0.01, ***p<0.001, trends 0.05≥# p<0.09; for comparisons between dependent groups: +p<0.05, ++p<0.01, +++p<0.001, trends 0.05 ≥ #’p<0.09.

The effect of QBP1 on the acquisition and consolidation of aversive memories was analyzed also by CFC **[Figure 3C]**, measuring the freezing time in the context where the fear conditioning was performed. The fear conditioning was well stablished as there were no significant differences between groups **[Figure 3C-i]** and both groups showed an increase of the freezing percentage throughout the test time [F_TRIAL(3,42)_=43.783 p<0.001 and F_INTERACTION(3,42)_=1.617 p=0.200; intersubject effect: F_TRIAL(1,14)_=2.738 p=0.120 and F_INTERACTION(1,14)_=91.541 p<0.001] **[Figure 3C-ii]**.

The **context test** was done 24 hours after the fear implantation. In order to analyze each minute of the context test, we referred the first minute as an adaptation time and we named the second to third minute as Min 1-2-3, so it could represent an analogy of the acquisition trials. We found that over the total test duration, WT mice showed a significant increase at the average freezing time in Min 1-2-3 *(x_M trial,* p=0.048) **[Figure 3D-i]**. Furthermore, there was an intrasubject effect of time and interaction [F_TRIAL(2,28)_=3.361 p=0.049 and F_INTERACTION(2,28)_=4.296 p=0.024], in which the inter-subject effect reflected that WT mice freezing behavior was significantly higher than those of the QBP1 mouse [F_TRIAL(1,14)_=4.709 p=0.048 and F_INTERACTION(1,14)_=95.460 p<0.001] **[Figure 3C-ii]**. Here QBP1 mice showed a significant decrease in the freezing percentage at min1 (p=0.005) and a trend in min2 (p=0.055) **[Figure 3C-ii]**, which suggests that QBP1 mice have not consolidated long-term fear conditioning. The **context test at 96h** showed no differences between both groups due to the freezing reduction at the control group [F_TRIAL(2,28)_=2.874 p=0.073 and F_INTERACTION(2,28)_=0.147 p=0.864; inter-subject effect: F_TRIAL(1,14)_=0.093 p=0.764 and F_INTERACTION(1,14)_=60.498 p<0.001] **[*Figure S3-A*]**.

In order to assess variations on anxiety levels after fear conditioning, an Elevated Plus Maze was performed on the analyzed mice after the context test 96h [***Figure S3-B*]**. In comparison with the anxiety levels showed in basal conditions by mice analyzed for hippocampal-dependent memories, QBP1 and WT mice significantly increased their times spent in closed arms after the fear conditioning. The selected aversive event (shock) was being stressful for both groups, which emphasizes that the differences observed in the CFC are due to a defect in long-term memory and not to differences in mouse anxiety.

An **extinction test** was also performed one week after fear implantation to study the relearning mouse abilities [***Figure S3-C*]**. Here they must avoid the learned fear as the same context is not aversive anymore. We found that only the WT mice showed a trend to change its behavior over time, which suggests that they are learning the new non-aversive context [F_TRIAL(3,42)_=5.151 p=0.004 and F_INTERACTION(3,42)_=0.334 p=0.801; inter-subject effect: F_TRIAL(1,14)_=3.872 p=0.069 and F_INTERACTION(1,14)_=99.155 p<0.001]. The **extinction context test (24h)** showed no differences between groups and none of them changed its behavior along time [F_TRIAL(2,28)_=0.314 p=0.733 and F_INTERACTION(2,28)_=2.534 p=0.097; inter-subject effects: F_TRIAL(1,14)_=0.392 p=0.542 and F_INTERACTION(1,14)_=65.102 p<0.001] [***Figure S3-D*]**.

A comparison between the freezing time on each context test was performed for QBP1 mice and control littermates **[Figure 3D]**. The results showed that QBP1 mice freezing values were constant along the different context tests [F_TOTAI TIME (2,20)_=1.502 p=0.249 and F_X_M_TRIAL (2,20)_=0.385 p=0.686], while WT mice significantly reduced their freezing times in correlation with the one showed at 24h after acquisition [F_TOTAI TIME (2,26)_=3.975 p=0.032 and F_X_M_TRIAL (2,26)_=4.371 p=0.024]. This final analysis reinforces the occurrence of an aversive memory impairment in the QBP1 mice.

### QBP1 mice after learning show altered posttranslational modifications of mCPEB3

Two weeks after the hippocampal-dependent memory tasks, the hippocampi of the tested mice were extracted and homogenized in order to analyze the mCPEB3 protein at each subcellular fraction. It has been shown that, after experiencing a learning process, mCPEB3 from these mice becomes oligomerized and increases its presence in the insoluble fractions^41^. As shown in the representative western blots of the performed extractions **[Figure 4A]**, the S2 fractions showed a main band of the mCPEB3 protein at 80 kDa, in close agreement with the molecular weight of the protein. The presence of a noteworthy higher molecular weight band (~120 kDa) suggested that mCPEB3 is also present with a post-translational modification^42^. The greater presence of this band in learned QBP1 mice (from now on, “learned” will refer to mice exposed to memorable events) could likely correspond to a higher SUMOylation of mCPEB3 and, therefore, to a greater presence of the inactive form of the protein. Other fractions were analyzed too, revealing slight differences between groups in learning conditions [***Figure S4-A*]**.

**Figure 4:**
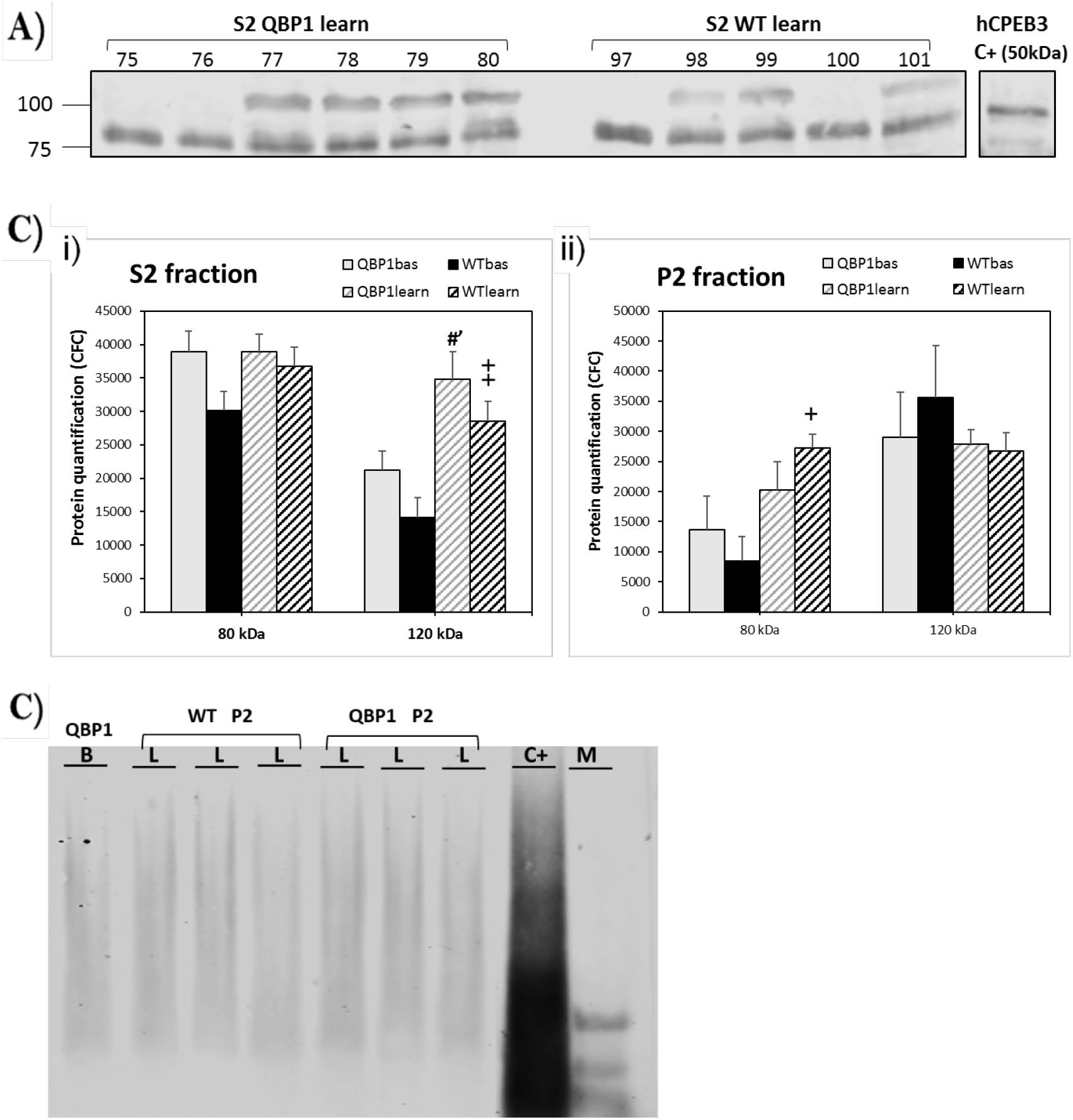
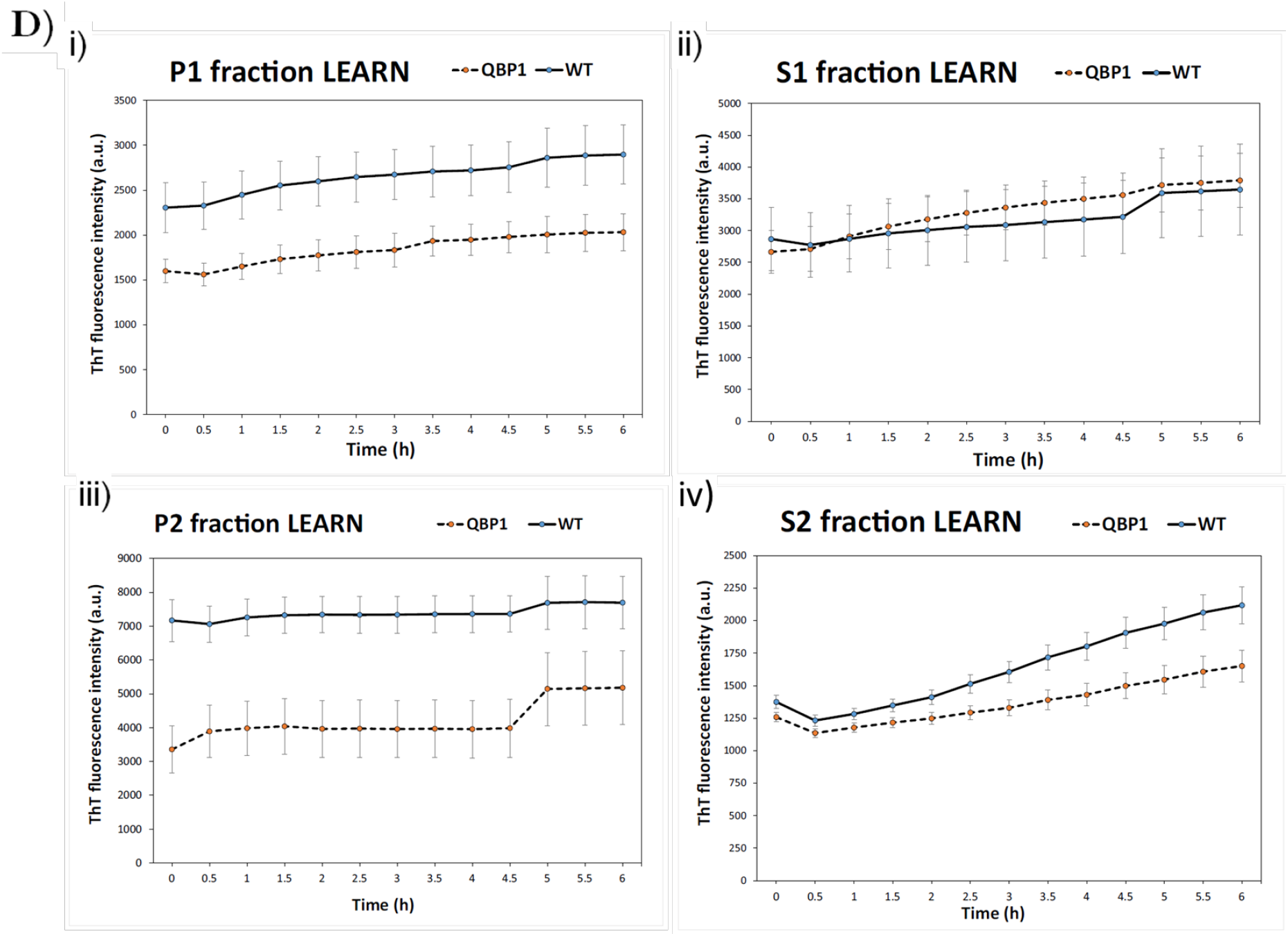
Analysis of mCPEB3 protein in the QBP1 transgenic mouse (learning *versus* basal conditions). **A) Western blot for S2 fraction (soluble) in learning conditions for mCPEB3**. QBP1 mice samples showed more intense posttranslational modification band than WT mice samples. Recombinant hCPEB3 (1-450aa) produced in *E. coli* was used as a positive control at 50 kDa of molecular weight, for both western blot and SDD-AGE experiments. **B) Quantification of western blot bands of S2 (i) and P2 (ii) fractions from mCPEB3**. There are no differences between the two groups, but in the soluble fraction the WT mouse significantly increases its 120 kDa band (p=0.003) while the QBP1 mice only showed a tendency (p=0.086). In the insoluble fraction **(ii)**, only the WT mouse significantly increases its values in the 80 kDa band (p=0.021), indicative of the greater oligomerization with learning. For comparisons between dependent groups: +p<0.05, ++p<0.01, +++p<0.001, trends 0.05 ≥ #’p < 0.09. **C) SDD-AGE for P2 fraction (insoluble) in learning and basal conditions for mCPEB3**. The oligomerized species can be observed in every sample analyzed, which suggest that the insoluble fraction contains mCPEB3 oligomers and there are no differences between groups. **D) Aggregation kinetics of Thioflavin-T fluorescence intensity of extracted hippocampi fractions in learning conditions**. The formation of mature amyloid with time is stronger for WT mice in all fractions, particularly in the insoluble fractions where there should be greater oligomerization (P1 and P2). Recombinant hCPEB3 (1-450aa) produced in *E.coli* was used as a positive control at 50 kDa of molecular weight, for both western blot and SDD-AGE experiments. Mean and SEM are represented, analyzing N=5 samples for each condition with three different repetitions of each one.

Furthermore, the quantification of the mCPEB3 bands in the wester blots showed significant differences in the soluble S2 and pellet P2 fractions **[Figure 4B, *Figure S5-A*]**. At the soluble fraction S2, there were no differences between both WT and QBP1 mice in the 80-kDa band but there was a tendency to increase for the QBP1 mice (p=0.086) and a significant increase for WT mice samples for the 120kDa band (p=0.003) **[Figure 4B-i]**. This difference could be due to the fact that it is in this fraction where the soluble protein is assumed to be in its active (80kDa) or inactive (120kDa, assuming SUMOylation) forms and, since the basal levels in QBP1 mice are already high for the 120 kDa band, the differences after the learning task are more difficult to evince. At the insoluble fraction P2 **[Figure 4B-ii]**, only WT learning samples significantly increased their values in correlation with basal conditions at the 80 kDa band. Thus, learning causes a greater presence of mCPEB3 at the insoluble fraction, suggesting a greater amyloidogenesis of this protein in WT mice in contrast with the case of QBP1 mice.

In order to analyze the amyloid properties of the extracted fractions, we performed **SDD-AGE** to study mCPEB3 oligomerization **[Figure 4C; *Figure S4-B]*** and a kinetics analysis using the **Thioflavin T (ThT) [Figure 4D]**, which is an amyloid stain specific for mature species. SDD-AGE results for the P2 fraction showed that the protein is found in the amyloid state while the aggregation kinetics results showed that in every insoluble fraction **[Figure 4D i-iv].** Samples from learned WT mice showed increased ThT fluorescence values. This suggested that in the hippocampi of the WT mice there is a greater amyloid oligomerization after learning than in the QBP1 mice. In contrast, the differences between groups in soluble fractions **[Figure 4D ii-iii]** were smaller and, specifically in S2, it seemed that the protein can aggregate along time, increasing its ThT values. These data were later confirmed through the increase of **Congo Red bound** over time,demonstrating that only the values of the S2 fraction of the “learn” WT mice increased significantly after incubation **[*Figure S4-C*]**.

Amyloid characterization in native conditions was performed by a **immunodot blot analysis**, measuring the immunoreactivity of the extracted samples in learning conditions with two conformational antibodies: A11, which is specific for prefibrillar oligomers and OC, which is specific for fibrillar oligomers and fibers^54^ [***Figure S4-D*]**. Although quantification showed no differences between QBP1 and WT “learn” samples regarding A11 and OC immunoreactivity [***Figure S4-E]***, the increased CPEB3 values for QBP1 samples seemed to indicate that mCPEB3 may be more accessible to interaction with the antibody in its native form. Furthermore, if we consider the high values of A11 immunoreactivity for the homogenate fraction, it seems that mCPEP3 is mostly found as a monomer in the QBP1 samples. Finally, it must be noted that our biochemical results confirm the involvement of the active amyloid form of CPEB3 in memory consolidation, whose role has been previously described^41,43^.

## Discussion

### Long-term hippocampal-dependent memories are impaired in the QBP1 mice

The QBP1 transgenic mouse has shown a significant impairment in memory consolidation, specifically after 24h of learning. This has been showed in both hippocampus-dependent memories (NOR, MWM) and aversive memories (CFC), which emphasizes that there is a general disruption of every new learned knowledge. Furthermore, this alteration is even present after the evening between two successive acquisition days, which is reflected in the increased IAR index of the QBP1 mice on the first days of MWM (worse performance in the first trial of each day). The impairment is specific for memory tasks, as everything seemed normal at activity cage, open field and, especially, in the 3-chamber test.

However, we have observed that in all tests QBP1 mice managed to resemble control mice behavior only by an intra-day repeated exposure or by its repetition over several days (from worse remembering at day 17 of MWM to a better remembering exactly the day after). Considering that CPEB3 oligomerization process is relatively quick^39^, although its expression is increased after a repeated exposure to the same stimulus^43^, in the QBP1 mouse we propose that the residual oligomerization would be increased with each exposure until it reaches a threshold for which its expression is enough to perform a normal memory consolidation.

In conclusion, it seems that the performance in the task gets worse with longer times after the last exposure, although the improvement is much greater if the exposure occurs on consecutive days: thus QBP1 mice have a long-term memory defect (<24h) and, at the same time, they are able to carry out some re-learning with the re-exposure to the task. This result is especially relevant because it means that the ability to learn remains intact and the only altered process by QBP1 is the persistence of long-term memory.

In order to better understand the molecular bases of the memory impairment that occurs in the QBP1 transgenic mouse, it is necessary to compare it with the knockout mice of the CPEB family described in the literature^32,42–44,55^. On one hand, as QBP1 is constitutively expressed and constrains the functional activation of CPEB3, the transgenic mouse acts as a global KO^55^ and shows a slightly better performance in the Morris Water Maze acquisition and normal anxiety at Elevated Plus Maze. On the other hand, the impaired consolidation of hippocampal-dependent and aversive memories may suggest a phenotype like that of the CPEB3 conditional KO (cKO) mice^43^. It must be noted that, the cKO of CPEB2 showed that the absence of this protein is also related to a long-term memory impairment^44^.

To explain how these different memory phenotypes could be consistent with the amyloidogenic properties of CPEB3^17,18,41^, we suggest that in the global KO, since CPEB3 is not present in the organism, it lacks the monomeric basal form whose action is repressive. Therefore,in the absence of repression, the process of memory consolidation is uncontrolled and gives rise to an enhancing memory phenotype, possibly driven by other protein or mechanism. However, in the case of the cKO, a basal CPEB3 concentration (repressor) is present so that the expression of new CPEB3 protein after synaptic stimulation is the step that is blocked. If the repressive basal form were the only species present, it might exert a strong action producing a phenotype of an impaired consolidation **[*Table S1*]**.

### Memory impairment in QBP1 mice correlates with a lowering of CPEB3 amyloidogenesis

We have investigated the underlying molecular mechanism of the alteration in the memory consolidation shown by QBP1 mice and found that it is concomitant with a lowering in the amyloidogenesis of CPEB3. So far, it has been shown that CPEB3 amyloidgenesis increases with learning and that CPEB3 KO mice have a long-term memory impairment^17,40–43,55^. However, with the introduction in the mice of the QBP1 peptide, to our knowledge, we are the first to directly modulate *in vivo* the amyloidogenesis of this functional amyloid, thus confirming its previously described involvement in memory consolidation and showing that it is not involved in learning acquisition.

Thus, we observed that the CPEB3 protein band (80kDa) only increases its intensity in the WT samples in the insoluble fraction after learning, as described in previous studies^43^.The ThT fluorescence intensity is also higher in the WT “learn” sample in this insoluble fraction, revealing that the increased band corresponds to a higher concentration of its amyloid species. This result confirms that QBP1 is blocking the oligomerization of CPEB3 reducing the function of its amyloid state, *i.e*. disrupting memory consolidation.

Furthermore, the presence of a greater ~120kDa band at the soluble fraction in QBP1 extracted samples suggests a likely posttranslational modification of CPEB3 by SUMO, a known repressor of CPEB3 amyloid state^42^. However, the quantification of this band only reflects a tendency to increase in learning conditions. Considering the lack of aggregation shown by ThT kinetics for “learn” QBP1 mice samples, we hypothesize that QBP1 may be favoring the monomeric form of the protein^50^, which would explain the existence of a greater band of SUMOylation. Thus, the band of ~120kDa seems to indicate an increased repression of CPEB3 by the expression of the peptide QBP1 in the transgenic mouse.

### A possible functional interplay between CPEB2 and CPEB3 in memory consolidation

To fully develop an explanatory model of the molecular mechanism involved in the QBP1 mouse phenotype, we must consider also the involvement of CPEB2 protein since the cKO of CPEB2 sheds light into the functionality of the different CPEB family of proteins. Interestingly, the conditional elimination of either CPEB2 or CPEB3 paralogs shows a similar phenotype, which suggests that both isoforms are being part of the same process and there is not redundancy. This also may explain the enhanced memory shown in CPEB3 constitutive KO^55^, assuming that CPEB2 protein may have a secondary role in memory consolidation.

Since the QBP1 blockade on CPEB3 is present since the transgenic mice are born, there may be a chance for some compensation so that the phenotype of the QBP1 mice is not as strong as that of the conditional KO^43^. We postulate CPEB2 as the main actor for this compensating mechanism. It is known that the increase in concentration of CPEB3 occurs mainly in dentate gyrus and CA3 only after stimulation^27^, while the functionality of CPEB2 has its main location in the neurons of CA1^44,56^. Furthermore, CPEB2 hippocampal expression is reduced even after synaptic stimulation^27^, while the levels of CPEB3 vary more significantly. We suggest that the oligomerization of CPEB3 is essential for the process since it regulates the establishment of longterm memory, while CPEB2 function would be more constant and stable over time. Thus, as the memory phenotypes of both QBP1 transgenic mice and the constitutive CPEB3 KO have showed^55^, CPEB2 could be in charge of part of the memory consolidation function compensating the absence of CPEB3 that occurs following the mouse’s birth.

Orb2 and *Ap*CPEB implication and regulation in different phases of memory consolidation have been studied in detail in the literature. In *Drosophila,* Orb2A isoform acts as a seed to induce the oligomerization of the constitutive Orb2B isoform^38^, thus forming essential heteroligomers for long-term memory. Likewise, in *Aplysia* it has been shown that the *Ap*CPEB4^57^ homolog cooperates with *Ap*CPEB^22^, so that both forms participate in long-term facilitation. Based on these evidences of the existence of an interplay between CPEB members of the same family of proteins in both species, we propose the existence of a similar mechanism in mouse. mCPEB2 is the likely candidate for an interaction with mCPEB3, based on the multiple analogies that can be found between Orb2A/Orb2B isoforms and mCPEB3/mCPEB2 paralogs regarding their function and regulation, which are summarized in **[*Table S2*]**.

### QBP1 as a potential drug for PTSD and ASD

Our results have shown the capability of the QBP1 peptide of inhibiting the oligomerization of CPEB3 protein in the QBP1 transgenic mouse, blocking in turn the consolidation of new memories in mammals. However, considering the polyvalence of QBP1, we cannot discard that the possibility that other amyloid/prion proteins involved in memory consolidation, like CPEB2, TIA-1, or huntingtin^18,44,58^, could be additional targets of QBP1. Likewise, pathological amyloidogenesis of neurotoxic proteins such as the polyQ tracts, *α*-synuclein or TDP43^49,50,59^, could also be inhibited by QBP1, which may be a rather beneficial side effect produced during this treatment if QBP1 derivatives were to be used to treat ASD or PTSD patients in the future (*see below*).

Previous studies have shown both that QBP1 is able to block the critical amyloidogenic conformational change at the monomer level of its target protein and that QBP1 blocks memory consolidation in a *Drosophila* model^39,49,50^ [***Figure S5-A*]**. However, the direct *in vivo* relationship between the memory blockade and the amyloidogenic state of the Orb2 protein has not been demonstrated yet. Our results in mice show the inhibitory effect of the QBP1 peptide on different types of memories (hippocampal-dependent and aversive memories) and provide a direct evidence of how amyloidogenesis can be modulated *in vivo,* explaining the long-term consolidation blockade by a concomitant reduction of mCPEB3 oligomerization [***Figure S5-B*]**.

Considering that QBP1 seems to be innocuous for mice and that it specifically impairs long term memory of new learned knowledge, including aversive memories, we propose QBP1 peptide as a promising lead compound for ASD and PTSD pharmacotherapy in humans. Thus, QBP1 derivatives could be used either to prevent the onset of ASD and PTSD (administrated after the trauma and before its consolidation) or as a therapeutic agent (combined with psychotherapy following a re-consolidation protocol) [***Figure S5-C*]**. It must be also noted that future preclinical studies should discard the possibility that QBP1 may also affect both anterograde and retrograde memories by the exogenous administration of QBP1, once the therapeutic time window of its administration is established. Remarkably, to the best of our knowledge, QBP1 peptide could be considered the first specific drug to treat mental disorders associated with the emotional valence of traumatic memories.

## Conclusions

Our study has established the proof-of-concept that the anti-amyloidogenic QBP1 peptide can block long-term memory consolidation in mice, extending its effectiveness to mammals. In addition to showing the harmlessness of the peptide through its constitutive expression, we have demonstrated how QBP1 affects the new knowledge acquired, with different memory effects depending on the tasker-exposing times on each task. Thus, as a lead compound for ASD and PTSD, it is quite remarkable that mice maintain the ability to learn while the drug blocks just the long-term maintenance of that knowledge.

## Materials and methods

### Production of the QBP1 construct

The insert expressing two copies of QBP1 (in tandem) fused to eGFP was constructed into the pCAGGS expression vector, which contains the CAG promoter into the rabbit *β*-globin splicing intron and ended with polyadenylation sites. The cDNA encoding QBP1(x2) was obtained from a plasmid isolated by phage display and used for protein expression in cell culture^48^, which was fused to the cDNA encoding EGFP. The sequence of all constructs was confirmed by DNA sequencing.

Then, the QBP1(x2)-EGFP cDNA was inserted in the p-EGFP-N1 vector as a template for its production, and then it was amplified to introduce the cDNA insert into the pCAGGS vector. The entire insert with the promoter and the encoding sequence of interest was excised with Sall and PstI from the expression vector. It was done following the similar protocols as previously described for other QBP1 constructions^48,60^.

### Production of QBP1 transgenic mouse

Fertilized eggs derived of a C57xC57 matting were extracted from the oviduct and were microinjected with the purified cDNA of the insert (enhancer-promoter-QBP1×2-EGFP-polyA terminator), by a subcontracted company (Japan SLC, Inc). A total of 200 injected ovules were implanted in pseudo-pregnant females which resulted in the birth of 3 lines of transgenic newborn mice. The incorporation of the transgene was examined by western blot in the brain lysate: brains were homogenized in 500 μL of TBS (pH 6.8) with protease inhibitor cocktail (Nacalai Tesque). About 10 strokes using a 2mL homogenizer was sufficient. The lysate was put in a 2 mL tube and 600 μL of 2 x SDS sample buffer was added using a cut tip. The lysate was pipetted about 5 times, boiled for 10 min, and cooled at room temperature. 5 μL of the lysate were mixed with 20 μL of SDS sample buffer and then 10 μL were used for SDS PAGE/Western blotting.

Furthermore, the constitutive expression was assured by genotyping mice from the purification of genomic DNA from mouse tail tissue *(Illustra tissue and cells genomic prep mini spin kit,* GE), with some modifications: incubation step with lysis buffer and proteinase K was carried out at 56°C for 3 hours and samples were eluted in 100 μL to improve concentration. Quantification of copy number was done by qPCR using the Taqman Copy Number assays protocol (Applied Biosystems), using Taqman probe versus GFP to detect the insert and Tert probe as a positive control.

### Animals

The experiments were carried out in male QBP1 transgenic mice of constitutive expression from 3 to 5 months of age and wild type littermates C57BL/6Ola (WT). The mice were generated from frozen embryos donated by Dr. Yoshitaka Nagai (previously frozen by the in-house facility of National Center of Neurology and Psychiatry), starting to analyze from the third homozygous generation, whose expression of the transgene was already stable. Frozen embryos were recovered by the Mouse Embryo Cryopreservation Facility at the Centro Nacional de Biotecnología-CSIC (Madrid). The mice were housed in groups of 4-6 per cage, under conditions of constant temperature (21-22 °C) and a light/dark cycle of 12 hours; with food and water *ad libitum.* The animals have been treated following the guidelines contained in RD 53/2013 and in accordance with European Community Guidelines (Directive 2010/63/EU), previously approved by the Bioethics Committee of the Cajal Institute.

Genotyping was carried out by the Molecular and Cellular Biology Unit of the Cajal Institute, based on mouse tail tissue (Illustra tissue and cells Genomic prep kit, GE Healthcare) and the copy number is quantified by quantitative PCR (qPCR, Taqman Copy Number assays, Applied Biosystems). A general protocol has been followed with a basic battery of behavior and neurological characterization, according to^61^, provided by the Unit of Animal Behaviour at the Cajal Institute. Weight, temperature and abnormal physical characteristics are weekly measured (balds, excessive grooming and integrity of the vibrissae).

### Statistical analysis

The statistical analysis was performed with IBM SPSS Statistics (Version 24.0, Armonk, NY, IBM Corp). The data were analyzed using an analysis of variance (one-way ANOVA or repeated measures), unpaired Student’s T test or Mann-Whitney U test (non-parametric cases); we previously checked the normality of the data by Shapiro-Wilk (N<50). For intrasubject differences, the Wilcoxon signed rank test (nonparametric cases) and the Student’s T test for related samples (parametric cases) were performed. Extreme values (outliers) were identified and visualized by boxplots using the aforementioned software and were removed from the analysis. The statistical significance was established at α=0,05. The data was handled in Excel software and, subsequently, graphs and images were created using GraphPad Prism 5 and CorelDRAW (v7). Values are always shown as mean ± SEM in tables and graphs.

### Hippocampi homogenization and fractionation

In order to analyze WT and QBP1 mice that have performed the hippocampal-dependent memory tasks, their hippocampi were homogenized using a detergent rich protocol^41,42^. The extracted hippocampi were incubated in 300 μL of lysis buffer (50 mM Tris pH 7.5, 1 mM EDTA, 10 mM KCl, 0.5% Triton, 0.5% NP40) for 1 hour in a cold room. Upon removing a small volume of homogenate for further analysis, the rest of the volume was centrifuged for 5 minutes at 4,000 rpm (Microfugue 18 centrifuge, Beckman Coulter) to obtain the cell debris fraction (P1) and a soluble fraction (S1), from which some volume was reserved too. This supernatant is ultracentrifugated at 65,000 rpm for 1 hour at 4°C in a TLA 100.1 rotor to obtain the soluble (S2) and insoluble (P2) protein fractions. Both pellet fractions were suspended in 50 μL of detergent buffer.

### Thioflavin-T fluorescence intensity (ThT): aggregation kinetics assay

The Thioflavin T (ThT) assay measures changes in fluorescence intensity of ThT after binding to mature amyloidogenic fibers^62^. The working solution of ThT (Sigma) was prepared at 1 mM in PBS buffer and a final reaction concentration of 50 μM was used. The assay was performed on Corning-384 plates, maintaining a final 20 μL volume: 1/6 dilution of the hippocampal extracts produced was analyzed. A continuous measurement was performed during 7 hours, programming the measurements every 120 seconds with a gain per well of 1300 at the FLUOStar OPTIMA Microplate Reader (BMG LABTECH), using two filters: a 440nm for excitation and 490nm for emission. Buffer with the same amount of ThT was used as a fluorescence background control.

### Western blot and SDD-AGE assay

Dissected hippocampi were homogenized in the detergent buffer and fractionated into soluble and insoluble fractions as mentioned. From each fraction, 10-20 μg were subjected to SDS-PAGE using 10% acrylamide gels and then transferred to a nitrocellulose membrane (0.45 μm, Amersham Protran) in tris-glycine buffer with 15% methanol. PBS 0.1% Tween-20 (PBS-T) with 5% non-fat-milk (Biorad) was used to block the membrane which was later incubated with the primary antibody: anti-CPEB3 rabbit polyclonal (ab10883) or anti-GAPDH rabbit polyclonal (ab37168). Washing steps were carried out with PBS-T 10% non-fat milk. IRDye 680LT Goat antirabbit (LI-COR) was used as secondary antibody and then revealed in Odyssey® CLx (LI-COR). The images obtained were analyzed by the Image J program in order to quantify protein level differences, in which two different western blots were analyzed for each sample (N=5 for each group).

The SDD-AGE assay is performed to visualize the presence of oligomerization in the fraction samples, maintaining native conditions. 1.5% agarose gel was prepared using TAE buffer with 0,1% SDS and protein samples were mixed with 5x loading buffer (5% glycerol, 0.1% bromophenol blue, 25mM EDTA) and a 2% SDS final concentration. Electrophoresis run for 2 hours at constant 70 V on ice. Proteins were transferred for 1.5 hours at constant 24V in TGS buffer with 15% methanol and 0.1% SDS final concentration. The antibody development process was carried out exactly as detailed for the western blot.

Extended information on additional experiments and detailed description of the experiments with mice has been included in the *Supplementary Material.*

## Acknowledgements

We thank Dr. D. V. Laurents for helpful discussions and English style suggestions and to Dr. Jose Ignacio Robles Sánchez for critical reading of the manuscript. We thank also the Molecular and Cellular Biology Unit (UBMC) from the Cajal Institute (CSIC) for the genotyping of mice samples. This work was supported by grants from the Spanish Ministry of Economy (SAF2013-49179-C2-1-R and SAF2016-76678-C2-1-R) to M.C.-V.

## Author Contributions

M.C.-V. conceived the project. P.L.-G, J.L.T, K.R.MG. and D.R.M. designed the animal experiments. P.L.-G, J.L.T, K.R.MG. analyzed the data and interpreted the results with the help of A.P.L. and A.S specifically with the 3-chamber and fear conditioning tests, respectively. H.A.P and Y.N. design the QBP1 construct and produce the QBP1 transgenic line, and performed the first characterization. D.R.M. recovered and maintained the transgenic line, characterizing the presence of the peptide in the hippocampus. P.L.-G designed the biochemical experiments, analyzed the data and interpreted the results. P.L.-G. wrote the first draft of the manuscript while M.C.-V. wrote the final draft.

