## Supplemental Material for "An anti-amyloidogenic treatment to specifically block the consolidation of traumatic events in mouse"

#### *Supplementary experimental data*

**Figure S1: Sociability-like behavior in the QBP1 transgenic mouse**.

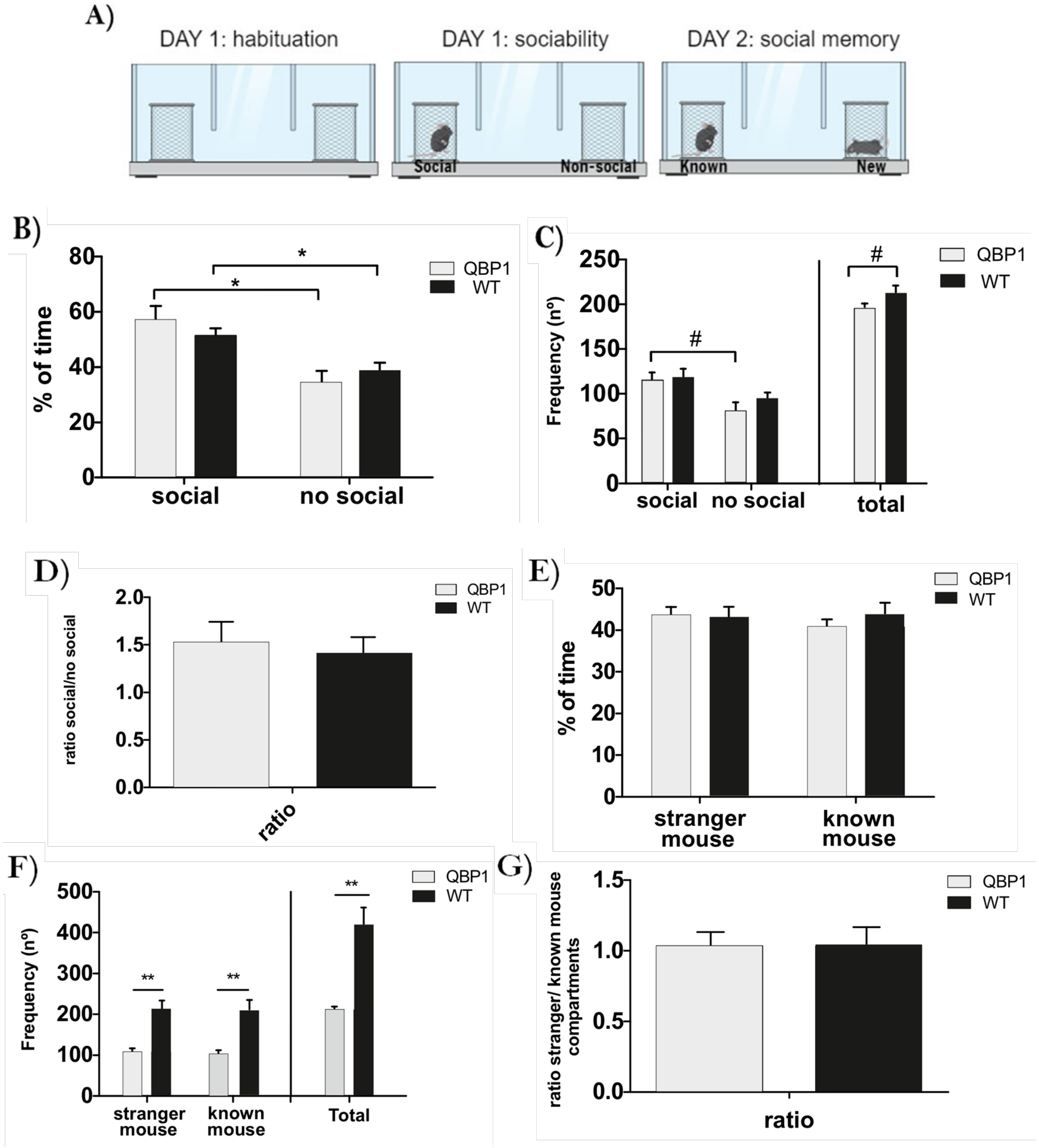

**Figure S2: Details of long-term memory impairment in the QBP1 mouse by the Morris water maze.**

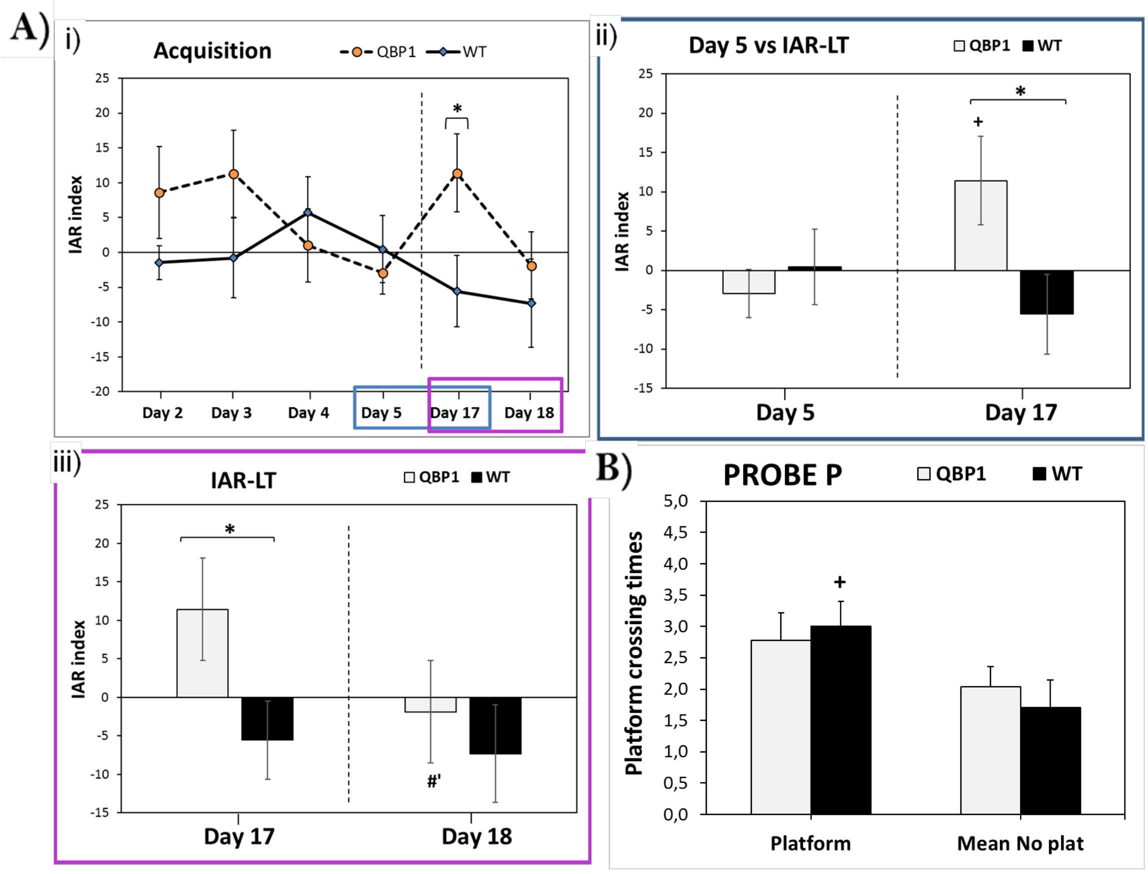

*Figure S3*: **Contextual fear conditioning confirmed the aversive memories impairment in QBP1 mice**

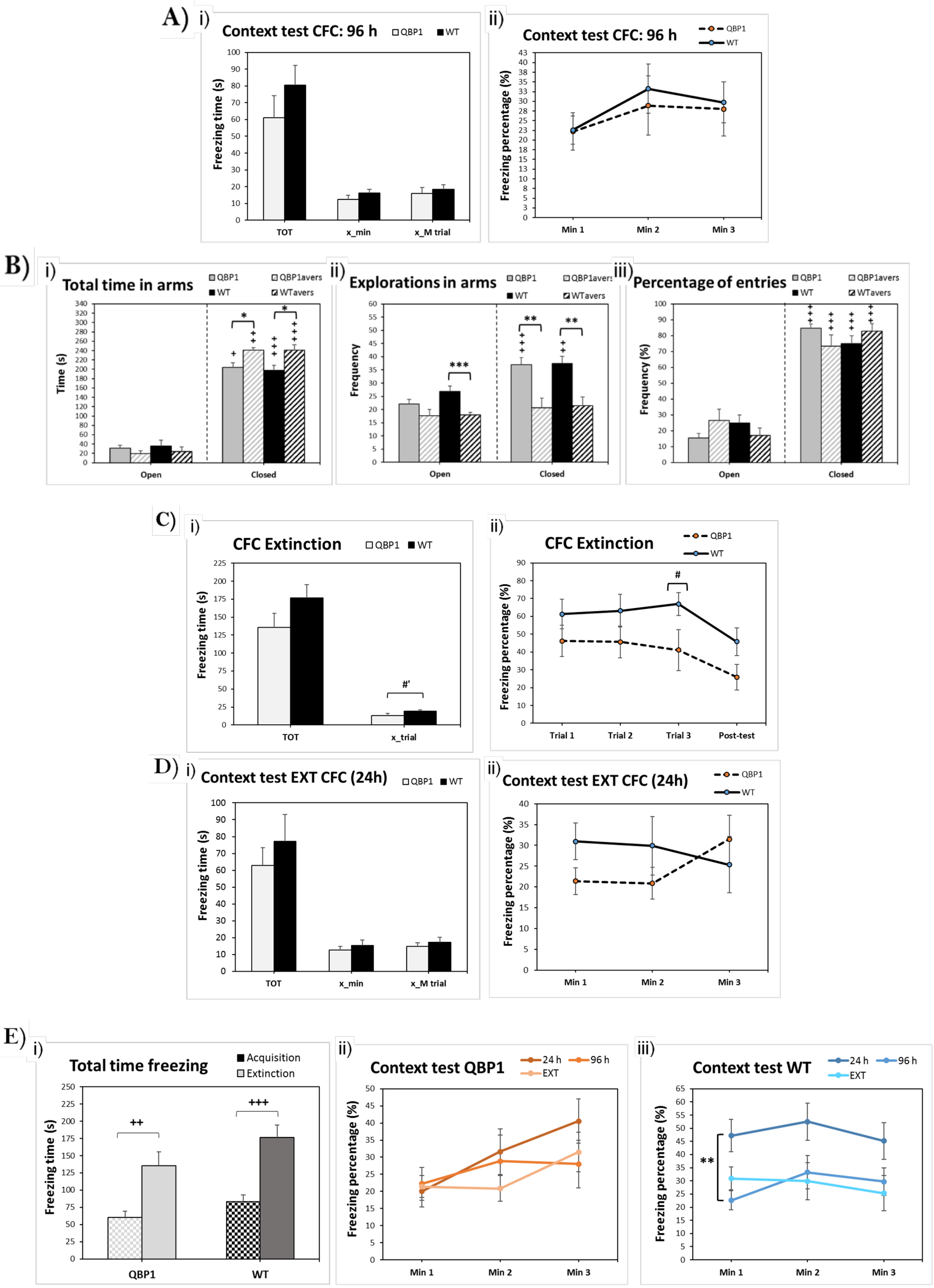

***Figure S4*: Additional biochemical studies showing differences among fractions and conditions.**

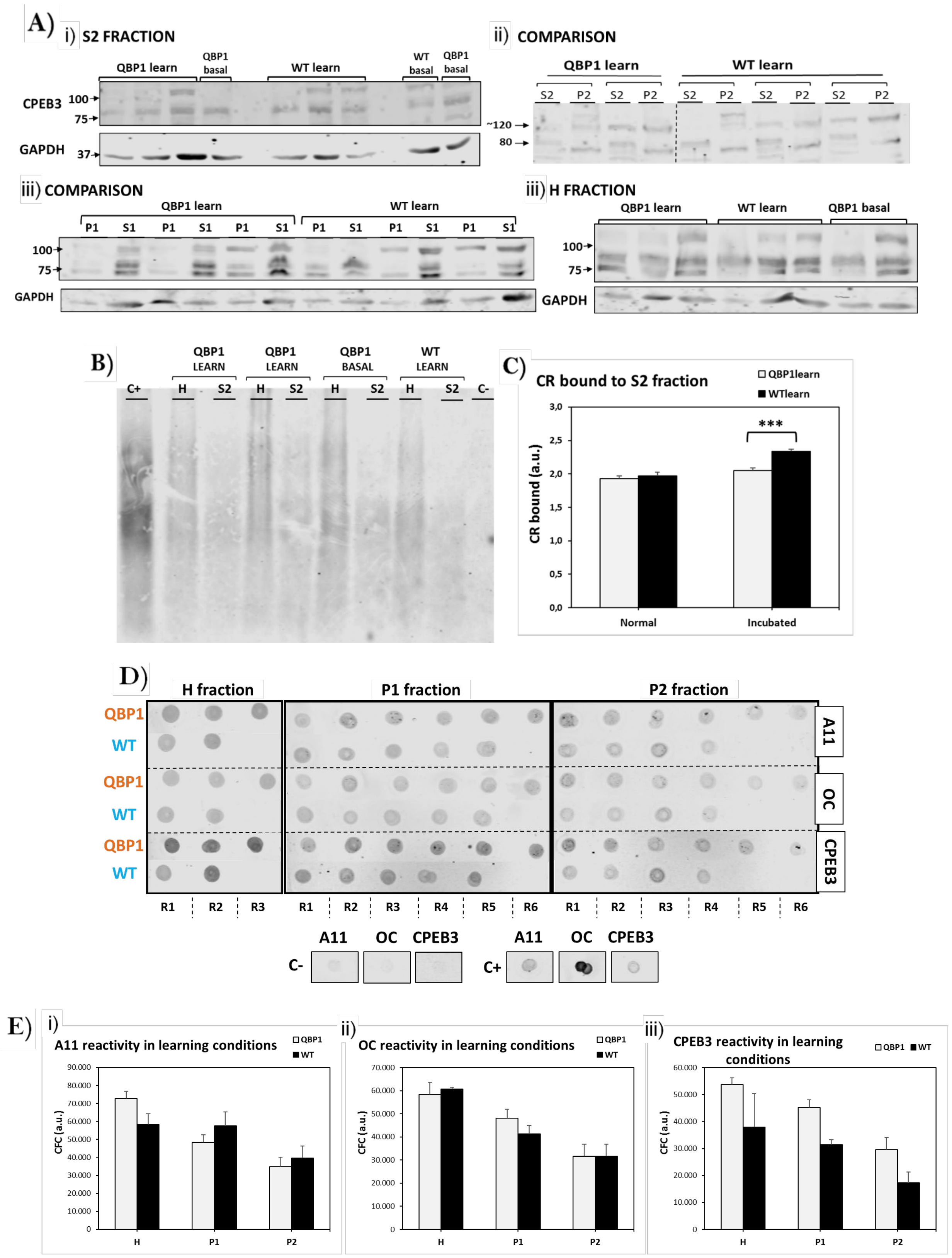

**Table S1: Review of CPEB KO literature in mice and a proposal for CPEB3-CPEB2 function in memory consolidation.** We have summarized the most representative data of the different mice analyzed in the literature that have studied the neuronal CPEB family. We establish a correlation between the memory phenotype and the hypothesis of the amyloidogenic state of CPEB3.

| AUTHOR | MICE | CPEB protein | CPEB3 state | PHENOTYPE | HYPOTHESIS |
|---|---|---|---|---|---|
| Chao (2013) | KO total | CPEB3 never is expressed<br>CPEB2 normal | ✘ Repressor<br>✘ Activator | **Enhanced** hippocampus-dependent memory and long-term spatial memory. | It could be that a **lack of inhibition from CPEB3 repressor to CPEB2 normal action** derived into an enhanced memory storage. |
| Fioriti (2015) | Conditional KO | CPEB3 expression is interrupted<br>CPEB2 normal | ✓ Repressor<br>✘ Activator | Mice have **impaired** long-term memory and LTP. | The same condition but in this case it did **exist CPEB3 repressor**, which might be strong enough to **block all memory** consolidation and CPEB2 function. |
| Lu (2017) | Conditional KO | CPEB3 is normal<br>CPEB2 expression is interrupted | ✓ Repressor<br>✓ Activator | Long-term memory consolidation is **impaired**. | If CPEB2 did not exist, CPEB3 in its both forms its not able to perform normal consolidation. **CPEB2 function is necessary for CPEB3 memory storage**. |
| Lu (2017) | Conditional KO rescue | CPEB3 is normal<br>No CPEB2 expression<br>CPEB2 target introduced in 31% of CA1 neurons | ✓ Repressor<br>✓ Activator | **Normal** long term-memory consolidation (phenotype rescued) | CPEB2 function is located in CA1 neurons and **CPEB3 (or its pathway) can act indirectly over its target** good enough to perform a normal memory consolidation. |
| QBP1 mouse | Transgenic | CPEB3 oligomerization is blocked<br>CPEB2 is normal | ✓ Repressor<br>✘ Activator | **Impaired** long-term memory: hippocampus-dependent and aversive memories. Oligomerization is reduced. | **QBP1 only acts over CPEB3** because if its acts over CPEB2 too the phenotype would be more severe than it is. **Maybe only CPEB3 shift its state to repressor to activator and thus allow memory consolidation regulating CPEB2 function**. |

**Table S2: Summary of functional analogies between neuronal CPEB systems: *Drosophila* Orb2 *vs.* mammalian CPEB family.** The correlation of functions, regulation mechanisms and proteins that interact with them is described, suggesting that in mammals a similar system may be present.

| ***Drosophila*** | **Mouse** |
|---|---|
| Orb2B protein is abundant and it is widely spread in the cell[1]. It has little oligomerization propensity and, if so, it is triggered by Orb2A[2]. | CPEB2 maintains a constant expression over time and independent of synaptic activity[3]. It maintains prion heritability but nothing is known about its amyloid characteristics[4]. |
| Orb2A has a low basal expression and is transcribed locally after synaptic activity[1,2]. It forms self-aggregates which show SDS-resistance[5]. | CPEB3 is barely expressed in basal conditions and it is only transcribed after synaptic activity and with temporal restriction[3]. Its amyloid and prion properties are largely described[6]. |
| Orb2A is regulated by different proteins due to the likelihood of being "*attacked*" by the proteasome. Those proteins are Tob and Lmk1[7,8]. | CPEB3 has a strict regulation over its amyloid conversion: SUMOylation is a constriction (SUMO-2)[9] while mono-ubiquitination is an enhancer (Neurl1)[10]. |
| Orb2A is essential for learning acquisition while Orb2B is needed for memory maintenance and persistence[11]. | It is yet unknown whether CPEB3 nor CPEB2 are implicated in early or late phases of memory consolidation. CPEB1 seems to be involved in extinction[12]. |
| Orb2A shows a short half-life in the oligomerized state[7]. | CPEB3 oligomers are short-lived, but it gets increased after new activity and persist longer times (recall and reconsolidation)[13]. |
| Orb2A/Orb2B can suppress its mRNA targets translation. Moreover, Tob interacts with Caf1/Pop2 in *Drosophila*[7]. | mRNA translation negative regulation: CPEB3 interacts with Tob1, which recruits Caf1 and promotes deadenylation[14] and repression. |

***Figure S5*: Summary of the findings on the effect of QBP1 peptide in the blockade of memory consolidation.**

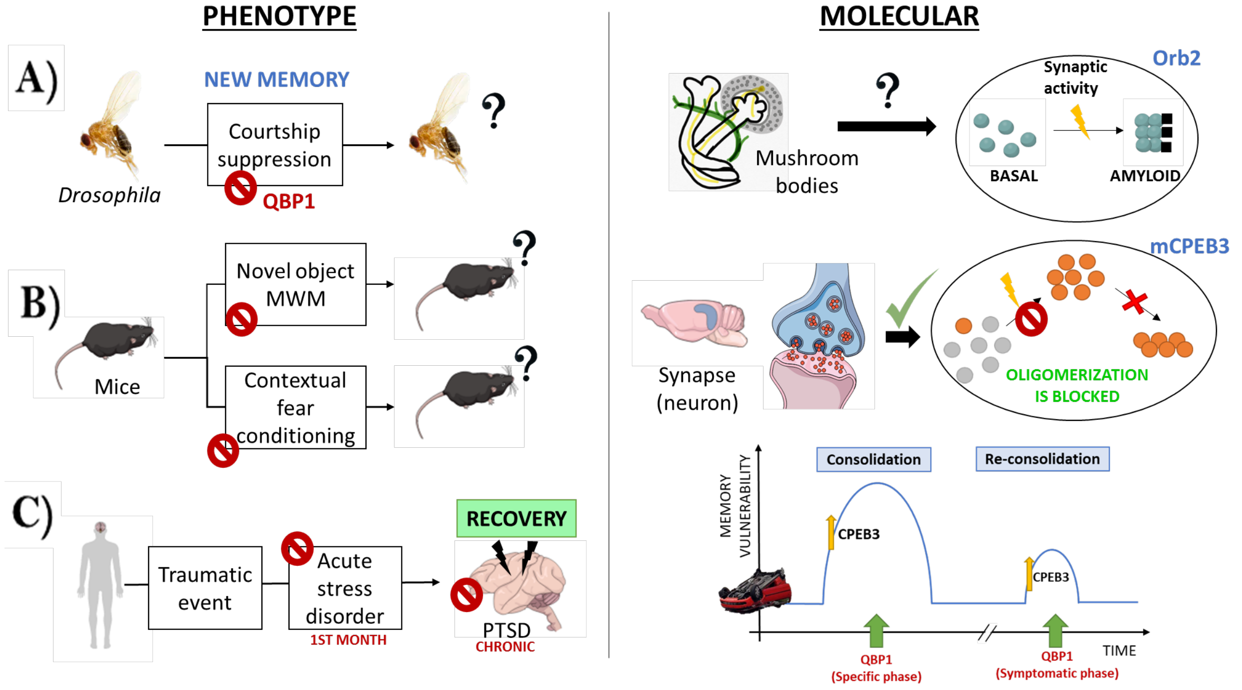


**Figure S1: Sociability-like behavior in the QBP1 transgenic mouse.**

**A)** Scheme of the two-day protocol in the 3-chamber test. After the habituation, the first day sociability-like behavior was analyzed and the second day social memory and novelty was assessed.

**B)** Percentage of total exploration time of each compartment. Both WT (p=0.033) and QBP1 (p=0.031) groups displayed normal sociability-like behavior by spending more time exploring the social compartment. No differences were found between groups neither in the social (p=0.303) nor the non-social compartment (p=0.436).

**C)** Number of entries in each compartment. WT mice entered equally both compartments (p=0.100) while QBP1 mice showed a trend for a higher frequency of entries to the social compartment (p=0.078). No differences were found between groups neither in the social (p=0.822) nor the non-social compartment (p=0.271).

**D)** The ratio of social preference showed any significant differences between groups. (T-test, p= 0.644). WT mice presented a trend for a higher total amount of exploration compared to QBP1 mice (T-test, p=0.056).

**E)** Percentage of total exploration time of each compartment on the second day of the test. No differences were found between groups in each compartment neither in the one with the stranger (p=0.809) nor in the one with a known mouse (p=0.411) and none of the groups (WT p=0.885, QBP1 p=0.424) showed preference for the stranger mouse.

**F)** Entry frequency on the second day. WT had a significant higher frequency than QBP1 mice exploring the known mouse (p=0.002), the stranger mouse (p=0.001) and a higher total frequency of entries (p=0.001). We did not found within-group effects in any of the compartments neither in WT (p=0.761) nor in QBP1 mice (p=0.773).

**G)** Ratio of novelty preference for a stranger mouse. No differences were found between groups (Mann-Whitney U test, p=0.087).

Data were analyzed by T-test and paired T test to assess inter- and intrasubject differences (showing mean ± SEM). For comparisons between independent groups: *p<0.05, **p<0.01, ***p<0.001, trends 0.05≥# p<0.09.

**Figure S2: Details of long-term memory impairment in the QBP1 mouse by the Morris water maze**.

**A)** IAR index of each day of test, reflecting the relationship between the first and the mean of all trials of each day. **i)** IAR index of WT mice (blue) is close to zero and then decreases as mice remember the platform location; for the QBP1 mice the index is higher the first days of acquisition and then decreases at day 5, but at day 17 it significantly increases (p=0.039) and 24h after it decreases again. **ii)** Comparison of the IAR index between day 5 of acquisition and day 17, which reflects an increase of the first trial difficulty at long term for QBP1 (p=0.042) with respect to the last day of acquisition. **iii)** comparison of the IAR index between day 17 and day 18, which reflects a loss of significance difference among groups and an improvement tendency in the QBP1 mice (p=0.091) 24h after the LTM test. A Friedman test was used to analyze differences between experimental groups in the IAR time course. A Mann-Whitney *U* was used to analyze differences between experimental groups in the IAR two-days comparisons, followed by a Wilcoxon signed-rank test to analyze intragroup differences between days.

**B)** Platform crossing times (probe P), in which only WT mice cross significantly more times the virtual platform location in the correct position than QBP1 mice instead of the same location in the other quadrants (p=0.017). A Mann-Whitney U test was used was used to analyze this comparison.

Data are showed as mean ± SEM. For comparisons between independent groups: *p<0.05, **p<0.01, ***p<0.001, trends 0.05≥# p<0.09.

***Figure S4*: Additional biochemical studies showing differences among fractions and conditions.**

**A)** Western blot that showed the comparison between S2 (soluble) and P2 (insoluble) fraction in learning conditions for mCPEB3. It suggests how both groups look similar and posttranslational bands are more present in the insoluble fractions

**B)** SDD-AGE to compare the oligomerization in H and S2 fraction (insoluble) in learning and basal conditions for mCPEB3. The signal is more intense for the homogenized fraction in every condition. It is hard to spot differences between groups, which means that other technique will be needed to quantify differences in the oligomerization.

**C)** Congo red bound in normal *versus* incubated (6h 37°C) condition is analyzed. CR is used as an alternative amyloid stain to quantify aggregation differences among conditions and groups. WT S2 sample is significantly increased when it is incubated (p<0.001), which suggests that the amyloid proteins involved are able to aggregate over time, while those in the QBP1 sample do not. The mean and the SEM are represented, analyzing N=3 samples for each condition. Data were analyzed by T test or Kruskal-Wallis to assess intra- and intersubject differences (showing mean ± SEM). For comparisons between independent groups: *p<0.05, **p<0.01, ***p<0.001, trends 0.05 ≥ #p < 0.09.

**D)** Immunodot blot analysis for homogenate(H) and insoluble fractions (P1 and P2) from extracted "learn" samples. In native conditions, every sample (R, mouse for each group) showed positive immunoreactivity for the A11 (prefibrillar oligomer conformation) and OC (fibrillar oligomer and fibril conformation) and anti-CPEB3 antibodies. Prefibrillar oligomers and fibrillar species of A*β*42 and the complete prion-like domain of hCPEB3 (1-450aa) recombinantly produced were used as positive controls.

**E)** Immunodot blot quantification for A11 **(i)**, OC **(ii)** and CPEB3 **(iii)** antibodies. There are no differences among QBP1 and WT learn samples at A11 and OC immunoreactivity, but QBP1 samples showed an increased CPEB3 value in the P1 fraction. H fraction was analyzed with N=3(QBP1)/2(WT) samples and P1/P2 fractions were analyzed with N=6(QBP1)/5(WT) samples.

***Figure S3*****: Contextual fear conditioning confirmed the aversive memories impairment in QBP1 mice**

**A)** Freezing behavior at different tests of the contextual fear conditioning. **i)** Total freezing time (TOT), freezing time average along minutes (x_min) and average freezing time of the analyzed minutes (x_M trial) are showed. There are no differences between groups. **ii)** Freezing percentage at context test after 96h of the acquisition evinced that there are no differences between groups and both behaved similarly (statistical values are defined in the main test).

**B)** Elevated plus maze results to compare anxiety levels in basal *versus* aversive conditions. **i)** Total time spent in arms showed that QBP1 and WT mice significantly increased their values after fear conditioning (p=0.028 and 0.016 in closed arms). Intra-subject differences showed that WT and QBP1 mice significantly increased its time in closed arms at both conditions (QBP1: p<0.001 for basal and p=0.027 for aversive condition; WT: p=0.000 for basal and p=0.005 for aversive condition). **ii)** Exploration frequency in arms is showed. WT and QBP1 mice significantly decreased their explorations in closed arms (p=0.003 and p=0.004, respectively) and only WT mice decreases theirs at open arms (p=0.001). Intra-subject differences showed that WT and QBP1 mice significantly increased their time in closed arms just at basal (p<0.001 for QBP1 and p=0.002 for WT). **iii)** Percentage of entries in arms is showed. There are no differences among groups. Intra-subject differences showed that WT and QBP1 mice significantly increased their percentage of entries in closed arms at both conditions (QBP1: p<0.001 for basal and p<0.001 for aversive condition; WT: p= 0.001 for basal and p<0.001 for aversive condition).

**C)** Extinction test of contextual fear conditioning, performed one week after the acquisition. **i)** Total freezing time (TOT) showed no differences between groups, but when the average freezing time in trials segments (x_trial) of the test is represented showed that WT mice trended to significantly increase their percentage in Trial 3 (p=0.090). **ii)** Freezing percentage along test time showed that WT mice freezing is higher than QBP1 mice, which is reflected by a trend in Trial 3 segment (p=0.056).

**D)** Context test 24h after extinction test. **i)** Total freezing time (TOT) and average freezing time along minutes (x_min) and average freezing time of the analyzed minutes (x_M trial) are showed. There are no differences between groups. **ii)** Freezing percentage at context test after extinction showed that there are no differences between groups and both behaved similar (statistical values are defined in the main test).

**E)** Freezing time intra-subject analysis at context test results for QBP1 mice and littermates. **i)** Total freezing time at acquisition and extinction of the fear conditioning is reflected for each group. QBP1 mice showed a significant increase on its freezing time (p=0.005) while WT mice reflected a very significant increase in the comparison (p<0.001). **ii)** QBP1 mice changes its behavior significantly time provided that 24 hours had passed since the test was performed [$F_{MIN\,(4,36)}$=2.137 p=0.096 and $F_{INTERACTION\,(2,36)}$= 10.636 p<0.001], while if the time is longer it remains without differences [inter-subject effect: $F_{MIN\,(2,18)}$=0.385 p=0.686 and $F_{INTERACTION\,(1,18)}$=86.025 p<0.001]. **iii)** Context test results for WT mice in each phase. WT mice always showed a stable behavior among time [$F_{MIN\,(2,48)}$=1.493 p=0.235 and $F_{INTERACTION\,(2,48)}$=0.787 p=0.539], but it could be found significant differences among phases in Min 1 (p=0.004). Their freezing time is higher 24h after the implantation and then it got lower, which is reflected by an inter-subject effect [$F_{MIN\,(2,24)}$=5.058 p=0.015 and $F_{INTERACTION\,(1,24)}$=145.800 p<0.001].

Data were analyzed by a mixed model ANOVA, Friedman test, or repeated measures ANOVA (for minute analysis), using T test, Mann Whitney *U*, and Wilcoxon signed-rank test to assess intra- and inter-subject differences (showing mean ± SEM). For comparisons between independent groups: *p<0.05, **p<0.01, ***p<0.001, trends 0.05 ≥ #p < 0.09; for comparisons between dependent groups: +p<0.05, ++p<0.01, +++p<0.001, trends 0.05 ≥ #'p < 0.09.

***Figure S5*: Summary of the findings on the effect of QBP1 peptide in the blockade of memory consolidation.**

**A)** Scheme corresponding to the experiments in *Drosophila*: at the phenotype level, only a single memory test (courtship suppression task) was used and the alteration of memories dependent on the Orb2 protein at the molecular level was determined[15].

**B)** Scheme corresponding to the experiments in QBP1 mouse: at the behavioral level, the blockage of hippocampal-dependent and aversive long-term memories has been demonstrated, as well as the harmlessness and effectiveness of the peptide in mammals. At the molecular level, direct evidence of the lowering of the oligomerization (orange) of the mCPEB3 protein has been established in the hippocampi of the learned QBP1 mice, thus confirming the effect of QBP1 on this functional amyloid *in vivo*.

**C)** Hypothesis of the expected effect of QBP1 in humans: the right dosage of the peptide would avoid the consolidation of traumatic events that could trigger PSTD and acute stress disorder. The administration is considered in a specific first phase (as a prophylactic: during the *first hours after the trauma*) and in a second symptomatic phase (as a therapy: in *reconsolidation processes*, as a reinforcement for psychotherapy).

### *Extended material and methods*

#### Activity cage and Open Field (OF)

The spontaneous motor activity of the animals ($N_{QBP1}$=8 and $N_{WT}$=10) was measured with an actimeter (VersaMax Legacy Open Field activity box, Omnitech Electronics), connected to a VersaMax analyzer and the VersaDat software. Two mice were placed *per* activity cage and the measurement was set up in five cycles of 1 minute, in a two-day protocol. Animals were habituated to the room for 30 minutes before the experiment and the cage was cleaned with ethanol before and between animals. Same protocol was used for Open field test, using one mice *per* cage and it was set up in two cycles of 5 minutes, done in a single day.

#### 3-chamber test

A 3-chambered apparatus with opaque external and Plexiglas internal walls was used to assess social preference and social memory/preference. On day 1, the social preference for a social *versus* a non-social compartment is tested. Both social and non-social compartments contain a wired cage placed in the middle of the chamber: an empty one corresponding to the non-social compartment, and a stranger mouse confined in the social chamber. On day 1, a habituation phase of the test takes place consisting of 5 minutes where the mouse is placed and confined in the central compartment, followed by 5 more minutes were the animal is allowed to freely explore the rest of the apparatus. Immediately afterwards, the animal is again confined in the central compartment for 1-2 minutes while the experimenter places a mouse in the wired cage of the social compartment. Consecutively, the animal is allowed to explore for 10 minutes the chamber and is videotracked. The time spent in every compartment of the chamber was calculated using Noldus Ethovision XT v11. On day 2 and 24 after the initial test, in order to assess novel preference for a new mouse, the subject mouse is allowed to explore the apparatus again where a stranger novel mouse has been placed in the previous empty compartment, and the known mouse of the previous test in the other one. Each test was counterbalanced within-groups to avoid biased results due to preferences for a side of the chamber.

#### Elevated plus maze (EPM)

The test apparatus consisted of four black polypropylene arms (Coulbourn Instruments, Whitehall, PA), with two "open" arms and two "closed" arms (walls of 30 cm height); all with 10 cm in width, 50 cm in length and at a height of 55 cm. At the beginning of each test, the animals were allowed to get used to the experimental room for 30 minutes and, at the beginning of the procedure they were placed in the center of the apparatus in front of an open arm. The animals were allowed to explore the apparatus for 5 min. The video signal was digitized and analyzed with the EthoVision XT 8.5 software (Noldus Information Technology, Leesburg, VA). An open arm entry was considered when the center of the mouse is in one of the open arms. The device was cleaned with ethanol before and between animals. It was performed in basal ($N_{QBP1}$=9 and $N_{WT}$=10) and after the fear implantation ($N_{QBP1}$=8 and $N_{WT}$=10).

#### Novel object recognition test (NOR)

NOR is a test that allows to analyze the discrimination of new objects with respect to objects already known, in a protocol of three phases: training, short-term test (both in the first day) and long-term test (second day) ($N_{QBP1}$=9 and $N_{WT}$=10). Protocol and guidelines followed were as described [16]. Mice were first habituated in the arena for 15 min and exposed to two different objects (A-B, training); 1 hour later short-term memory was tested for 10 min, in which mice are exposed to A-C objects. 24 hours later, long-term memory is tested (10 min) and mice are exposed to A-D objects. The mice were recorded using a video camera and the exploration time was counted using EthoVision XT 8.5 software: the observer who blindly analyzed the results considered the active approach of the mouse nose within a one centimeter range of the object as an exploration of the object. The discrimination index (DI) was determined by the difference in exploration time expressed as a ratio of the total time spent exploring the two objects: good discrimination is settle at 0,20 DI value, considering that there must be significant differences between the training and the testing phases of the test.

#### Morris Water Maze (MWM)

The experimental mice were tested ($N_{QBP1}$=9 and $N_{WT}$=10) based on the standard procedures of the Animal Behavior Unit at the Cajal Institute (CSIC) based on well stablished protocols [17]. The test is based on the ability of the mouse to learn the location of a platform placed in one of four quadrants of a cylindrical pool. Finding the platform is the only way out, so the mouse must reduce the escape latency within successive trials, in the first days of acquisition. After a habituation test (day 0) in which the preferences between quadrants in the different experimental groups were discarded, the animals learned to find a hidden platform during the following 5 days, through 4 trials/day (60 seconds each test, plus 20 seconds on the platform). If an animal cannot reach the platform, the experimenter places it on top of it. Subsequently, the animals were subjected to a trial on the seventh day, without the platform, to assess the preference for the quadrant of the platform (Probe). During this probe, path tracks were analyzed to measure the time spent swimming in the platform quadrant (probe Q) compared to the time in the other three quadrants, or the time spent swimming above the virtual position of the platform (probe P), compared to the time above the virtual mirror positions or the platform in the other three quadrants. In days 17 and 18, animals were exposed to a new series of four trials a day for two days, recording the escape latency.

#### Contextual Fear conditioning for aversive memories (CFC)

Fear conditioning was implanted in QBP1 transgenic mice (N=8) and WT littermates (N=10) using the UGO BASILE ANY-maze controlled Fear Conditioning System (mouse cage). In the acquisition day, after the adaptation time (180 seconds) mice were exposed to three trials of 30 seconds: a footshock is used as the conditioned stimulus (CS, 0.8 mA of intensity) and it is given in the last 2s of the trial, in which an unconditioned stimulus is present (UCS, 30s of continuous white sound -70 dB-). Each trial was spaced by a fixed intertrial time (ITI: 50s) and then 30s are recorded after the last ITI. 24 and 96 hours after the acquisition, context test was assessed: mice were returned to the conditioning chamber and freezing behavior was measured during 5 minutes (no shock and 250 lux). The design of the experiment and the subsequent collection of the data has been done with its behavioral tracking program (ANY-maze software). Extinction was performed seven days after fear acquisition using the same protocol used in the fear implantation, although CS was removed.

In the data analysis, TOT measure reflected the total freezing time and *x_min* is the average freezing *per* minute at context test (in which "x" reflected the symbol for the average). As acquisition and extinction comprises three phases (adaptation, trials and post-test), each context test was also divided into test segments: first minute is adaptation and the final minute is post-test, thus calling Min 1-2-3 to the central minutes, simulating what the different trials would be. It was called *x_trial* to the average freezing time in trials segments of acquisition and extinction test; whereas *x_M trial* is the average freezing time along those central minutes in context test (Min 1-2-3).

#### Congo-red binding

For the spectroscopic test with CR, a working solution at 60 µM in buffer (5 mM potassium phosphate, 150 mM NaCl, pH 7.4) was prepared and filtered immediately before use. 5 µL of protein (10 µM, incubated until the corresponding time in each case) was added to 5 µL of the CR solution, so that a final ratio of 1:2 (protein:CR) is maintained. The UV-visible spectrum is recorded in the Nanodrop between 400 and 700 nm, collecting the absorbance values at 480 and 540 nm to subsequently calculate the concentration of CR bound [18,19].

#### Dot blot assay

This technique was used to detect, analyze the conformation and identify proteins in its native state immobilized in a nitrocellulose membrane by their binding to antibodies that recognize specific structural conformations: A11 antibody, which detects the existence of prefibrillar (toxic) oligomers, and OC antibody, which detects fibrillar oligomers and mature fibers. Extracted hippocampal samples where diluted 1/6 and then 2 µl were placed on a 0.45 µm nitrocellulose membrane (Amersham Protran). The membrane was blocked using 10% fat-free milk (Blotting-Grade Blocker non-fat dry milk, BIO-RAD) and subsequently the membrane was incubated with the corresponding primary antibody: A11 1:2000 antibody (Invitrogen, ref: AHB0052) and OC antibody 1:2500 dilutions (Millipore, AB2286). Finally, it was incubated with the LI-COR antibodies as described for the western blots. Pre-fibrillar oligomers and fibrillar species of Aβ42 and the complete prion-like domain of hCPEB3 (1-450aa) produced *in vitro* were used as positive controls.

### *References for supplementary material*

10. Pavlopoulos, E. *et al.* Neuralized1 activates CPEB3: A function for nonproteolytic ubiquitin in synaptic

plasticity and memory storage. *Cell* **147**, 1369–1383 (2011).
